# Interpretable Peripheral Blood Cell Classification via Zero-Shot Vision-Language Concept Bottleneck and Soft Decision Tree

**DOI:** 10.64898/2026.07.14.738462

**Authors:** Kai Chen, Ting Hu

**Affiliations:** School of Computing, Queen’s University, Kingston, ON, Canada

**Keywords:** interpretable machine learning, concept bottleneck model, vision-language model, soft decision tree, peripheral blood cell classification, hematology

## Abstract

**Motivation:** Deep learning classifiers for medical image analysis typically function as black boxes, disclosing neither the image features underlying their predictions nor the reasoning by which individual decisions are reached. Peripheral blood cell classification exemplifies this challenge: experienced laboratory professionals identify cell types through structured morphological criteria—nucleus shape, chromatin texture, nucleus-to-cytoplasm ratio, granularity, and staining properties—yet existing automated systems cannot express their reasoning in these same terms, impeding clinical audit and verification.

**Results:** We present a two-stage interpretable pipeline that addresses both levels of opacity. In the first stage, a frozen domain-adapted vision-language model (ConceptCLIP) projects each cell image onto a 70-dimensional vector of morphological concept scores via zero-shot cosine similarity, eliminating the need for per-image concept annotations. In the second stage, a Soft Decision Tree (SDT) classifies cells solely on these concept scores, producing a deterministic, concept-based decision path for each prediction. On BloodMNIST (eight cell types, 3,421 test images), the full pipeline achieves 94.86% test accuracy—approximately 3 percentage points below the black-box ceiling—while providing fully traceable decision logic. Post-training histological annotation confirms that the learned routing logic aligns with established hematological morphology criteria and reveals an emergent separation of immature granulocyte subtypes (promyelocyte versus metamyelocyte) without subtype supervision, demonstrating that concept-based decision trees can recover clinically meaningful distinctions beyond the granularity of the training labels.

**Availability and implementation:** The source code, trained SDT weights, precomputed concept score data, and inference scripts are publicly available at https://github.com/aquamarineaqua/CLIP-CBM-SoftDecisionTree.

## 1. Introduction

Deep learning models have achieved remarkable performance in medical image classification tasks, often matching or exceeding trained specialists [1, 2, 3]. However, their opacity poses a fundamental barrier to clinical deployment: a model that cannot provide an explanation to its predictions is difficult for clinicians to audit, verify, and trust. Peripheral blood cell classification— the task of identifying cell types from microscopic images of blood smears—exemplifies this challenge. Experienced laboratory professionals classify cells by recognising a structured set of morphological features including: nucleus shape, chromatin texture, nucleus-to-cytoplasm (N:C) ratio, granularity, and staining properties. A reliable automated system would not only predict the correct cell type but also express its reasoning in these same morphological terms, enabling clinical audit and verification.

Existing explainable AI (XAI) approaches in medical imaging are predominantly post-hoc: they explain already-trained black-box models after the fact. Methods such as Grad-CAM [4] highlight spatial regions of influence; SHAP [5] and LIME [6] assign importance scores to input dimensions. While informative, these methods answer only *where* the model attends, not *what* morphological features it perceives or *how* it integrates them into a diagnosis—the questions most clinically relevant for diagnostic support systems.

Concept Bottleneck Models (CBMs) [7] address this limitation by replacing opaque latent features with predefined, semantically meaningful concepts, making the feature representation interpretable by construction. However, traditional CBMs require per-image concept annotations during training— an impractical demand in medical domains where labelling effort per concept per sample is prohibitive. CLIP [8], a vision-language model trained on 400 million image–text pairs via contrastive learning, provides an annotation-free alternative: concept scores can be computed as cosine similarities between image embeddings and text-based concept prompts, without any per-image labelling. This principle underlies recent annotation-free CBMs in natural image domains [9, 10]; we extend it to blood cell morphology using ConceptCLIP [11], a CLIP variant pretrained on medical image–text pairs with emphasis on clinical concept-level alignment.

A CLIP-based concept bottleneck renders the feature representation interpretable, but a downstream MLP or SVM remains a black box. To achieve end-to-end transparency, we replace the conventional classifier with a Soft Decision Tree (SDT) [12]. The SDT is a differentiable model that routes each input through a tree of learned oblique splits; at inference time, a hard routing mode yields a deterministic, sequential decision trace that mirrors the coarse-to-fine reasoning of clinical differential diagnosis.

We make the following contributions, validated on the BloodMNIST benchmark:

1. We demonstrate that a zero-shot CLIP-based concept bottleneck with 70 domain-informed morphological concepts retains ≈98% of the discriminative information in the full image embedding, incurring approximately 1.9 percentage points of accuracy loss relative to the black-box model.
2. We show that replacing the downstream classifier with a Soft Decision Tree incurs only an additional ≈1.3 percentage point accuracy cost, yielding a fully transparent pipeline at a total cost of ≈3 percentage points below the performance ceiling.
3. We provide histological analysis of the learned decision tree, demonstrating alignment with established hematological morphology criteria and revealing an emergent separation of immature granulocyte subtypes (promyelocyte versus metamyelocyte) without supervision.

The proposed two-stage pipeline is illustrated in Figure 1.

**Figure 1.**
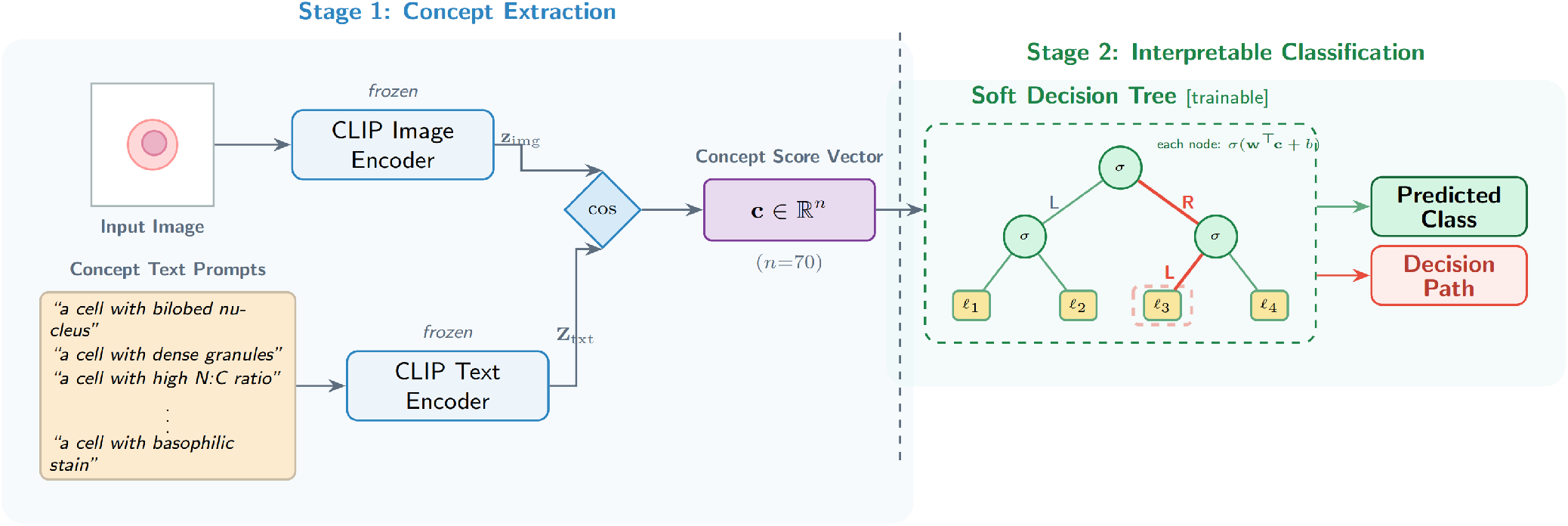
Overview of the proposed pipeline. Stage 1: a frozen ConceptCLIP backbone converts each blood cell image into a 70-dimensional morphological concept score vector via zero-shot cosine similarity with text-described concepts. Stage 2: a Soft Decision Tree classifies cells based solely on these concept scores, producing a deterministic, concept-based decision path for each prediction.

## 2. Methods

### 2.1. Dataset

All experiments in this study use BloodMNIST [13], a bench-mark from the MedMNIST collection comprising microscopic images of individual peripheral blood cells derived from the dataset of Acevedo *et al*. [14]. We use the 224×224-pixel, three-channel version to preserve fine-grained morphological detail. The dataset contains eight classes of normal peripheral blood cells: basophil, eosinophil, erythroblast, immature granulocyte (comprising promyelocytes, myelocytes, and metamyelocytes), lymphocyte, monocyte, neutrophil, and platelet (Figure 2). The original MedMNIST training set (11,959 images) and validation set (1,712 images) are merged to form a single training set of 13,671 images; the official test set of 3,421 images is held out for all evaluations to ensure comparability with published benchmarks.

**Figure 2.**
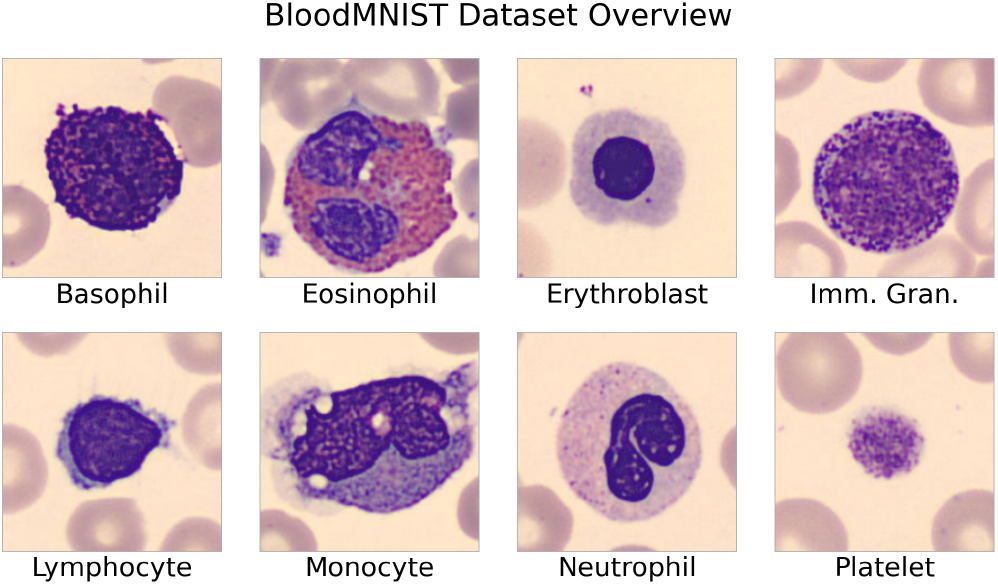
Sample images from BloodMNIST (May–Grünwald–Giemsa staining, 224*×*224 px). Eight classes are shown: basophil, eosinophil, erythroblast, immature granulocyte, lymphocyte, monocyte, neutrophil, and platelet.

### 2.2. Concept Set Design

The concept set is derived from hematological morphology literature and covers five categories: (i) *nuclear morphology and N:C ratio* (shape descriptors including segmented, band, bilobed, reniform, and round nucleus; chromatin texture; nucleoli; N:C ratio); (ii) *cytoplasmic tone and texture* (basophilic/eosinophilic staining, vacuoles, perinuclear hof); (iii) *cytoplasmic granules and inclusions* (azurophilic, eosinophilic, and basophilic granules; toxic granulation; pathological inclusions such as Auer rods); (iv) *non-leukocyte blood elements* (erythrocyte and platelet morphology visible in the field); and (v) *preparation and technical artifacts* (stain precipitate, motion blur).

Four concept sets of varying granularity (number of concepts: *n* = 30, 50, 70, 90) were designed across these categories; the optimal size was determined empirically (Section 3, Table 1). The full 70-concept listing is provided in Supplementary Table S1.

**Table 1.** Classification accuracy as a function of concept set granularity on the BloodMNIST test set.

| Concept Set Size ($n$) | Classifier | Test Accuracy |
| --- | --- | --- |
| 30 | Logistic Regression | 0.9225 |
| 50 | Logistic Regression | 0.9436 |
| 70 | Logistic Regression | 0.9567 |
| 90 | Logistic Regression | 0.9594 |

Each concept is encoded as an ensemble of ten text prompt templates (e.g., “a blood cell image indicating {}” and “a micrograph showing the feature of {}”; full list in Supplementary Note S2). The ten text embeddings per concept are averaged to produce a single robust representation, mitigating the linguistic bias of any single phrasing.

### 2.3. CLIP-Based Concept Score Extraction

Given the frozen ConceptCLIP backbone, for each input image the image encoder produces an embedding **z**_img_ ∈ ℝ^*D*^. For each of the *n* concepts, the ensemble-averaged text embedding 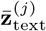is computed by averaging the ten prompt embeddings. The concept score for concept *j* is:

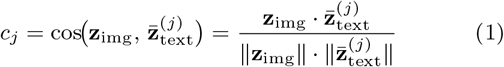

The concatenation of all *n* scores yields the concept score vector **c** ∈ ℝ^*n*^, which replaces the raw image embedding as input to all downstream classifiers. The text prompts function as natural language annotations, and the VLM’s cross-modal alignment provides the concept-scoring bridge, bypassing per-image concept labelling.

### 2.4. Soft Decision Tree Architecture and Training

#### Architecture

An SDT of depth *d* contains 2^*d*^ − 1 internal nodes and 2^*d*^ leaf nodes. Each internal node *i* computes a routing probability:

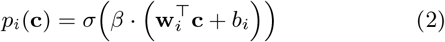

where **w**_*i*_ ∈ ℝ^*n*^ and *b*_*i*_ are learnable parameters, *σ*(·) is the sigmoid function, and *β* is an inverse temperature. Because **w**_*i*_ is dense over all *n* concept dimensions, each node performs an oblique (hyperplane) split—a weighted combination of multiple concept scores—providing robustness to noisy individual scores. The probability of sample **c** reaching leaf *l* is:

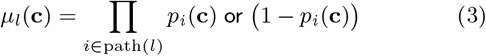

and the final prediction is *P* (*y* | **c**) = Σ_*l*_ *µ*_*l*_(**c**) · *Q*_*l*_, where *Q*_*l*_ is the learned class distribution at leaf *l*. At test time, hard routing assigns each sample to a single leaf via argmax over leaf-reaching probabilities, yielding a deterministic decision path.

#### Depth selection

Depth *d* = 4 (15 internal nodes, 16 leaf nodes) was selected empirically: *d* = 3 plateaued at ≈92% accuracy, while *d* = 5 provided no accuracy gain over *d* = 4 but doubled the leaf count, degrading interpretability.

#### Training

The SDT is trained on the merged 13,671-image training set. The loss function combines cross-entropy on the soft mixture output with a balance penalty *C* = −*λ* · 2^*−k*^ · [log(*α*_*i*_) +log(1 − *α*_*i*_)], where *α*_*i*_ is the average left-routing probability at node *i* at depth *k* and *λ* is a regularisation coefficient; the penalty encourages balanced tree utilisation by penalising routing probabilities far from 0.5. Training uses Adam [15] with learning rate 5 ×10^*−*3^, weight decay 5 ×10^*−*4^, *β* = 2, and *λ* = 5 ×10^*−*5^. The ConceptCLIP backbone is frozen throughout.

### 2.5. Pruning and Tree Annotation

Following training, a post-hoc pruning and annotation procedure extracts interpretable decision logic. A node or leaf is pruned if: (1) fewer than 50 training samples are routed through it under hard inference; or (2) for leaf nodes, the maximum predicted class confidence is below 80%. These criteria identify statistically unsupported or diffuse components.

Retained nodes are annotated by analysing routing behaviour at each internal node. Within the subset of training samples routed to a given node under hard inference, the top-*K* samples (*K* = 50) with the highest decision scores **w**^*⊤*^**c** + *b*— routed left—and the bottom-*K* with the lowest scores—routed right—are visualised and examined through histological expertise. Because each dimension of **c** corresponds to a named morphological concept, the signed weight vector **w** assigns each concept a directional contribution to the routing decision; the extreme-scoring images serve as visual archetypes of the two branches’ morphological character.

## 3. Results

### 3.1. Concept Set Granularity

To determine the optimal concept set size, we evaluated the four concept sets using logistic regression as the downstream classifier (Table 1). Logistic regression was selected to isolate the effect of concept set composition from classifier complexity. Accuracy increases monotonically with concept dimensionality: +2.1 pp (percentage points) from *n* = 30 to 50, +1.3 pp from 50 to 70, but only +0.27 pp from 70 to 90, indicating saturation. The 70-concept set is adopted for all subsequent experiments as the optimal balance of coverage and parsimony.

### 3.2. Accuracy–Interpretability Progression

#### Black-box upper bound

Using the full ConceptCLIP image embedding as input, a single-hidden-layer MLP achieves 0.9798 test accuracy (Table 2), establishing the performance ceiling.

**Table 2.** Black-box upper bound: classification accuracy using the full ConceptCLIP image embedding.

| Features | Classifier | Test Accuracy |
| --- | --- | --- |
| Image embedding | MLP (1 hidden layer) | 0.9798 |
| Image embedding | XGBoost | 0.9678 |
| Image embedding | Logistic Regression | 0.9386 |
| Image embedding | SVM | 0.9582 |

#### Concept bottleneck

Replacing the image embedding with the 70-dimensional concept score vector, the MLP and SVM each achieve 0.9611— approximately 1.9 percentage points below the ceiling (Table 3). This confirms that the 70-concept bottleneck retains the large majority of discriminative information in the full embedding, and attests to ConceptCLIP’s cross-modal alignment quality.

**Table 3.** Concept bottleneck performance using the 70-dimensional concept score vector. XGBoost underperforms relative to linear classifiers, consistent with axis-aligned splits being sensitive to correlated and noisy concept scores.

| Features | Classifier | Test Accuracy |
| --- | --- | --- |
| Concept scores ($n=70$) | MLP (1 hidden layer) | 0.9611 |
| Concept scores ($n=70$) | XGBoost | 0.9216 |
| Concept scores ($n=70$) | Logistic Regression | 0.9567 |
| Concept scores ($n=70$) | SVM | 0.9611 |

#### Full pipeline

The SDT achieves 0.9486 under hard inference, incurring an additional ≈1.3 pp over the best concept-bottleneck classifier— the cost of the tree’s structural constraint. Table 4 summarises the complete accuracy–interpretability progression.

**Table 4.** Summary of the accuracy–interpretability trade-off. The proposed full pipeline (bold) achieves complete decision-logic interpretability at ≈3.1 pp total accuracy cost, decomposing into ≈1.9 pp from the concept bottleneck and ≈1.3 pp from the SDT structural constraint. Concept scores are labelled “partially” interpretable because individual scores are cosine similarities rather than calibrated concept-presence probabilities.

| Stage | Features | Classifier | Accuracy | Interp. Features? | Interp. Logic? |
| --- | --- | --- | --- | --- | --- |
| Black-box ceiling | Image embedding | MLP | 0.9798 | No | No |
|  | Image embedding | XGBoost | 0.9678 | No | No |
| Concept bottleneck | Concept scores ($n=70$) | MLP | 0.9611 | Partially | No |
| | Concept scores ($n=70$) | LR | 0.9567 | Partially | No |
| | Concept scores ($n=70$) | SVM | 0.9611 | Partially | No |
| **Full pipeline** | **Concept scores ($n=70$)** | **SDT ($d=4$)** | **0.9486** | **Partially** | **Yes** |

### 3.3. Decision Tree Structure and Biological Findings

#### Pruning outcome

Applying the pruning criteria (Section 2.5, Pruning and Tree Annotation), 7 of 16 leaf nodes are pruned: L1 (34 samples), L5 (53), L6 (1), L7 (0), L10 (16), L11 (7), and L15 (29), collectively accounting for ≈1% of the training set. The remaining 9 active leaves are retained for annotation. Two internal nodes (IN10, IN12) have both children pruned and are excluded from interpretation. The class distribution across all 16 leaf nodes is shown in Figure 3; every cell type is covered by at least one active leaf, and 8 of 9 active leaves achieve 99% dominant-class confidence.

**Figure 3.**
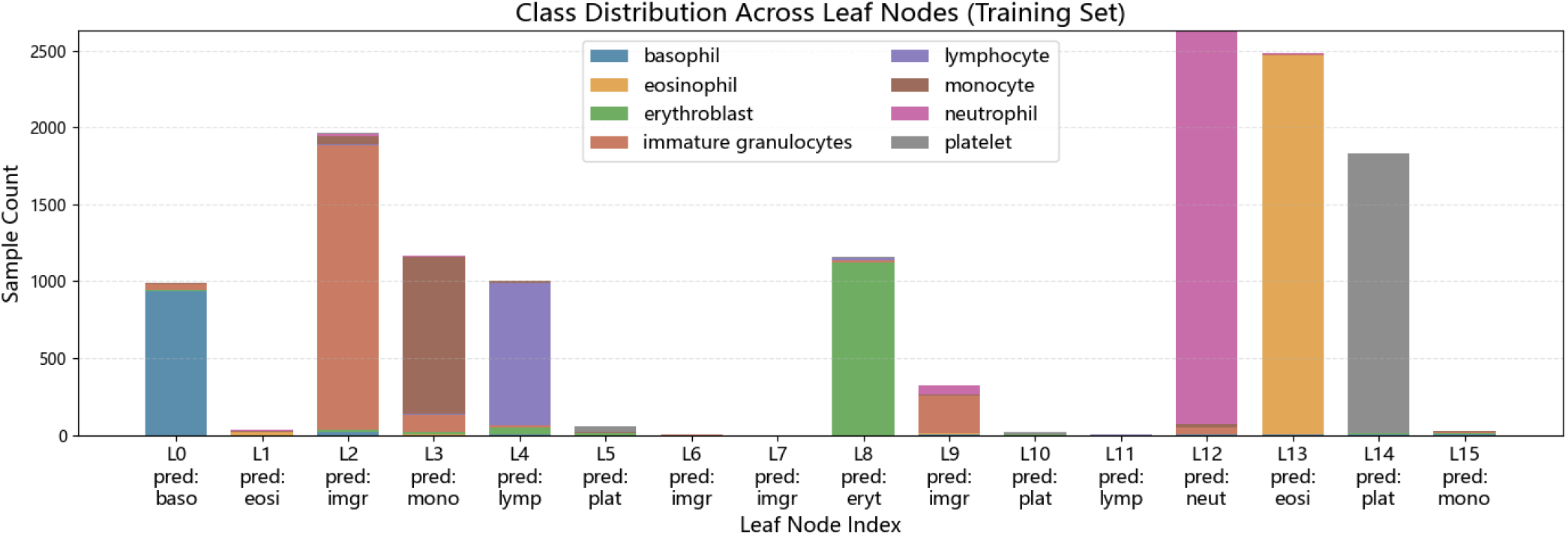
Class distribution across the 16 leaf nodes of the trained SDT on the training set. Active leaves are labelled with their predicted class; pruned leaves receive negligible sample volume.

#### Hierarchical decision logic

The tree’s routing logic follows a coarse-to-fine organisation (Figure 4). The root node (IN0) partitions the dataset based on cytoplasmic staining tonality under May–Grünwald–Giemsa staining: cells with predominantly basophilic (blue/violet) cytoplasm are routed left, while cells with eosinophilic (pink/red) cytoplasm or small cell bodies are routed right. This chromatic dichotomy is the primary organising axis in routine blood smear analysis.

**Figure 4.**
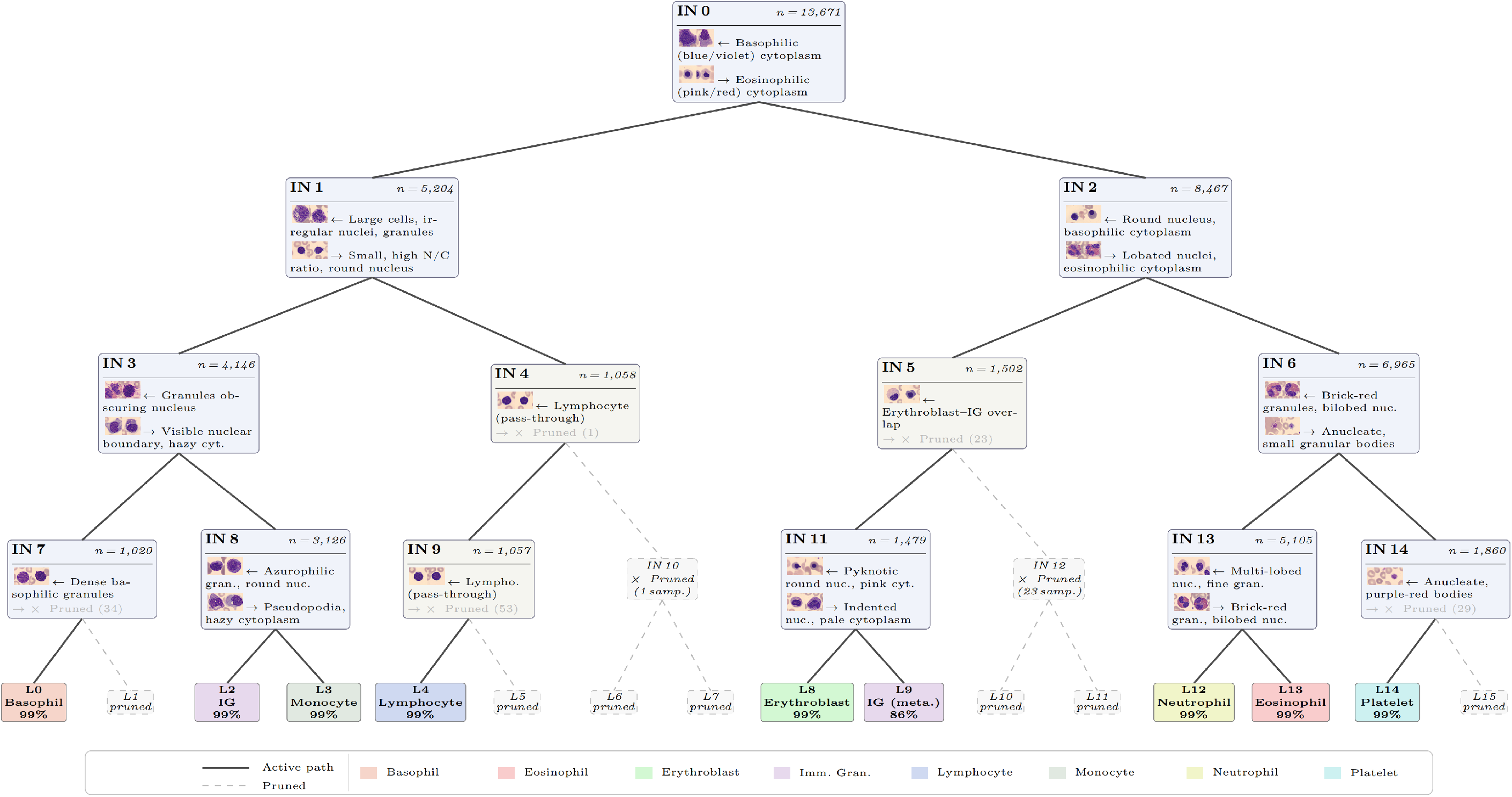
Annotated structure of the trained depth-4 Soft Decision Tree after pruning. At each internal node, the learned oblique split partitions samples based on a linear combination of the 70 concept scores. Representative training-set images are shown for the high-routing-score (left branch) and low-routing-score (right branch) populations, with brief histological characterisations. Pruned subtrees (dashed lines) correspond to leaves with fewer than 50 training samples or prediction confidence below 80%.

At layer 2, IN1 separates lymphocytes from remaining basophilic-toned cells using small cell body, high N/C ratio, and round dense nucleus—precisely the standard criteria for lymphocyte identification. IN2 isolates erythroblasts from the eosinophilic-toned population by detecting round nuclei with dense basophilic-to-polychromatic staining. Deeper layers resolve within-group distinctions: IN3 separates basophils from immature granulocytes and monocytes via granule density and degree to which granules obscure the nucleus; IN8 discriminates immature granulocytes from monocytes based on granule type and nuclear contour regularity; IN6 separates platelets from granulocytes; and IN13 distinguishes eosinophils from neutrophils by granule characteristics and nuclear lobation. Complete per-node weight heatmaps and representative images are provided in Supplementary Note S1.

#### Emergent immature granulocyte subtype separation

Immature granulocytes are captured by two distinct decision paths leading to separate leaf nodes: L2 (via IN0 → IN1 → IN3 → IN8) and L9 (via IN0 → IN2 → IN5 → IN11). Inspection of representative samples reveals morphologically distinct populations (Figure 5).

**Figure 5.**
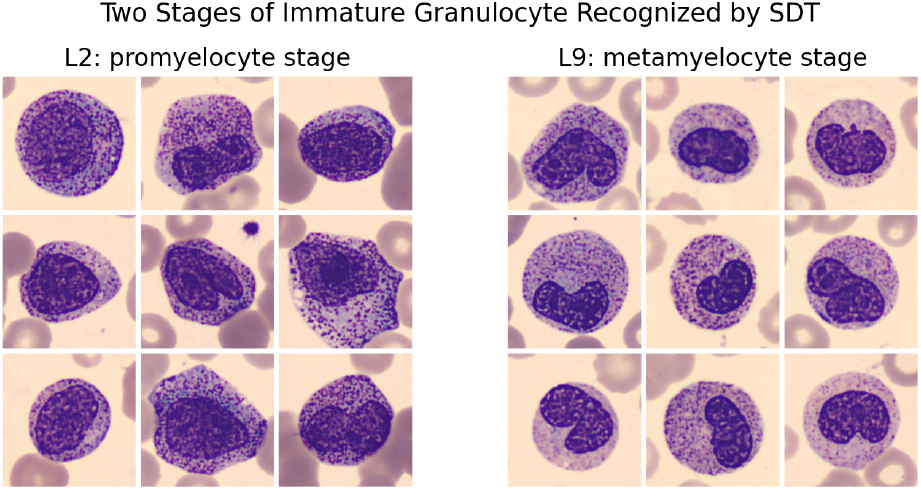
Morphological comparison of immature granulocyte subtypes captured by distinct decision paths in the SDT. Top row: representative samples from L2 (promyelocyte-like features — round nucleus, dense azurophilic granules). Bottom row: representative samples from L9 (metamyelocyte-like features — indented nucleus, paler cytoplasm). No subtype supervision was provided during training.

The IG samples reaching L2 exhibit features consistent with the promyelocyte stage: round-to-oval nuclei without indentation, dense azurophilic (primary) granules, and a relatively high N/C ratio. Their cytoplasmic staining character aligns these cells with the basophilic subtree. The IG samples reaching L9, by contrast, exhibit features consistent with the metamyelocyte stage: indented (reniform) nuclei, paler pink cytoplasm approaching the appearance of mature neutrophils, and a reduced N/C ratio, placing them in the eosinophilic subtree. Blood MNIST provides only a single “immature granulocyte” label encompassing all maturation stages; no subtype annotations were available during training.

## 4. Discussion

### 4.1. Clinical Relevance of the Learned Decision Logic

The layer-by-layer annotation of the SDT reveals a routing structure that closely mirrors the coarse-to-fine strategy employed in clinical differential diagnosis of peripheral blood cells. The root split on cytoplasmic staining tonality, followed by progressive refinement based on nuclear morphology, granule characteristics, and cell size, reproduces the sequence in which morphological features are hierarchically evaluated in standard hematological practice. That this structure emerged without any supervision on the decision sequence—only cell type labels were provided during training—demonstrates that concept-based SDTs can recover domain-consistent reasoning from distributional patterns in the concept score space.

The emergent separation of immature granulocyte subtypes illustrates this capacity for intra-class structure discovery. Although promyelocytes and metamyelocytes share a single class label in BloodMNIST, the SDT partitioned them into distinct leaf nodes driven purely by morphological gradients in the 70-concept vector. This observation also provides retrospective justification for tree depth *d* = 4: a depth-3 tree with eight leaves would enforce a one-to-one leaf-to-class mapping, precluding such intra-class discovery. More broadly, morphological heterogeneity within a single diagnostic label is the norm for pathological categories such as blast cells, atypical lymphocytes, and reactive forms in clinical hematology. Concept-based decision trees, by providing representational redundancy through extra leaves, may be applicable to unsupervised subtype discovery in settings where fine-grained labels are unavailable but morphological distinctions are clinically consequential.

### 4.2. Concept Projection as Semantic Dimensionality Reduction

The concept bottleneck is algebraically a linear projection: because CLIP embeddings are L2-normalised, cosine similarity reduces to a dot product, and the concept score vector results from projecting the image embedding onto the concept-defined subspace. Unlike classical dimensionality reduction (PCA, t-SNE [16], UMAP [17])—which derive projection axes from data statistics—this projection is onto semantically labelled directions specified by human domain knowledge and mediated by the VLM’s cross-modal alignment (Supplementary Figure S3). Each dimension of the resulting vector has a name and a domain-grounded interpretation.

This distinction bears on downstream interpretability: when classical dimensionality reduction feeds a decision tree, splits reference anonymous axes; when the concept projection feeds the SDT, each split operates on a named linear combination of morphological features. The ≈1.9 pp accuracy cost of the concept bottleneck quantifies the gap between the full perceptual representation (what the VLM encodes) and the cognitive representation (what can be named in domain language): features that are discriminative but orthogonal to all 70 concepts are discarded by construction, because they cannot be expressed in the morphological vocabulary and therefore cannot be audited by a clinician.

### 4.3. Limitations

Several limitations should be noted. First, the 70-concept set is manually curated; there is no formal guarantee of completeness or absence of redundancy. Individual concept scores are cosine similarities rather than calibrated probabilities of concept presence, and are subject to cross-concept correlation. The SDT’s oblique splits mitigate this by aggregating multiple concepts, but per-weight interpretations should be treated as directional rather than precise.

Second, all experiments were conducted on BloodMNIST, a balanced dataset of eight typical cell types. The dataset does not include pathological variants (blast cells, atypical lymphocytes, reactive forms) or rare subtypes critical in clinical practice. Generalisability to complex morphological taxonomies, class imbalance, or out-of-distribution morphologies remains to be established.

Third, quantitative results are reported from a single training run; run-to-run variance due to random initialisation is not quantified, and the reported accuracy should be interpreted as a representative rather than an expected value.

Finally, scalability is a structural limitation: at depth 4, 16 leaves are inspectable by a human annotator, but exponential leaf growth at greater depths would degrade practical interpretability for tasks with substantially more classes or finer morphological distinctions.

## 5. Conclusion

This work presented an interpretable pipeline for peripheral blood cell classification that achieves transparency at two levels: the feature level, via a 70-concept CLIP-based bottleneck, and the decision level, via a Soft Decision Tree. On BloodMNIST, the full pipeline retains 94.86% test accuracy at a total cost of approximately 3 percentage points relative to the black-box ceiling, decomposing into ≈1.9 pp from the concept bottleneck and ≈1.3 pp from the SDT structural constraint. Post-training annotation confirmed that the learned routing logic aligns with established hematological morphology criteria and revealed an emergent separation of immature granulocyte subtypes without subtype supervision.

The concept bottleneck performs a deliberate transition from a perceptual representation to a cognitive one: the discarded information is precisely that which cannot be expressed in the morphological vocabulary and therefore cannot be audited by a clinician. Whether the resulting accuracy cost is acceptable is context-dependent—screening settings may prefer the black-box ceiling, while diagnostic settings requiring clinician verification or regulatory transparency may favour the full pipeline.

Several directions remain open. The 70-concept set is manually curated; automated concept discovery—via large language models or data-driven methods—could complement or validate manual design. Evaluation on datasets with pathological variants, class imbalance, and finer-grained labels is necessary to establish clinical applicability. Scalability beyond depth 4 will require methods for selective depth expansion or hierarchical tree ensembles to maintain interpretability as the number of classes grows. Finally, multi-seed evaluation should be conducted to quantify run-to-run variance and distinguish methodological stability from properties of a single checkpoint.

## Author Contributions

K.C. conceived the study, designed the concept set, implemented the full pipeline, conducted all experiments, performed the histological analysis, and wrote the manuscript. T.H. supervised the project, provided conceptual guidance, and critically revised the manuscript. All authors read and approved the final manuscript.

## Data and Code Availability

BloodMNIST is publicly available as part of the MedMNIST collection (https://medmnist.com/) [13]. The source code, trained SDT weights, and inference scripts are publicly available at https://github.com/aquamarineaqua/CLIP-CBM-SoftDecisionTree.

