## Supplementary Material for "Interpretable Peripheral Blood Cell Classification via Zero-Shot Vision-Language Concept Bottleneck and Soft Decision Tree"

#### Contents

- Supplementary Figures S1–S3
- Supplementary Tables S1–S2 (Concept Set Compositions)
- Supplementary Note S1: Complete SDT Node Annotations
- Supplementary Note S2: Text Prompt Templates

#### Supplementary Figures

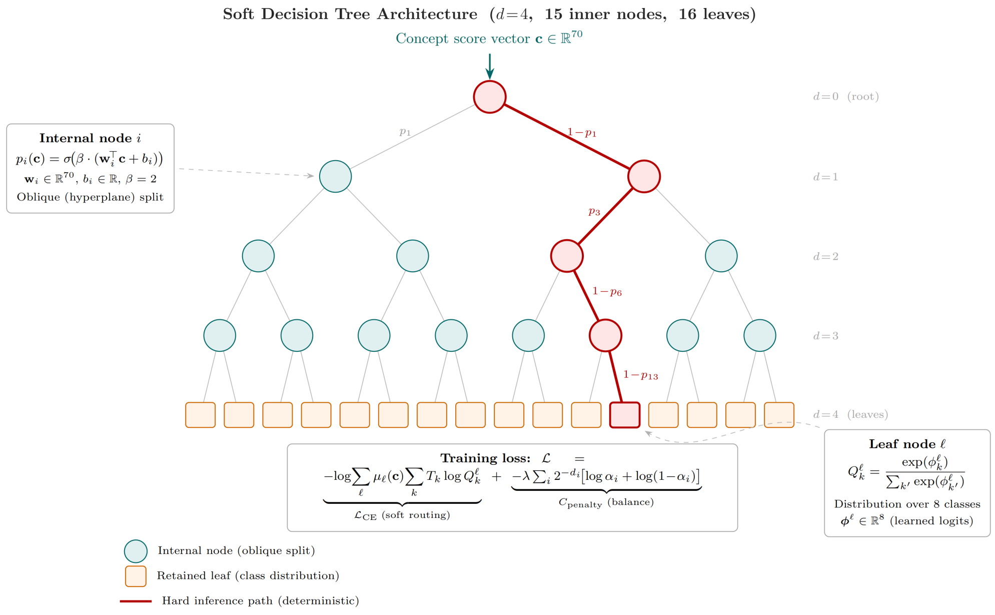


**Figure S1** Architecture of the Soft Decision Tree (SDT). Each internal node $i$ computes a routing probability $p_i(\mathbf{c}) = \sigma(\beta \cdot (\mathbf{w}_i^\top \mathbf{c} + b_i))$, directing samples left (high score) or right (low score). During training, all leaves receive gradient signal via soft routing; at inference time, hard routing assigns each sample to a single leaf via argmax, producing a deterministic decision path.

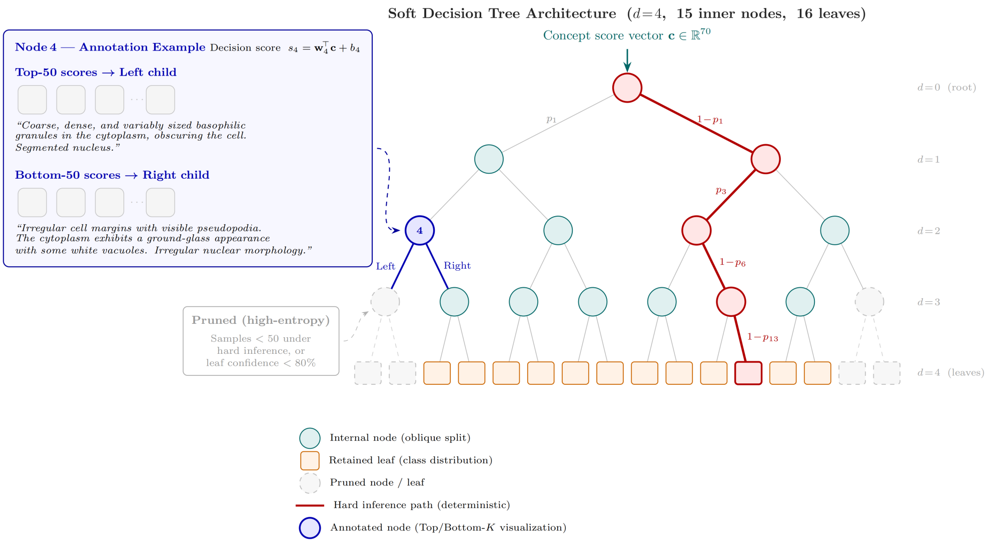


**Figure S2** SDT pruning and annotation procedure. Following training, nodes or leaves with fewer than 50 routed training samples or leaf prediction confidence below 80% are pruned. Retained nodes are annotated by visualising the Top-50 (highest decision score, left branch) and Bottom-50 (lowest decision score, right branch) training samples, which serve as morphological archetypes for the two routing directions.

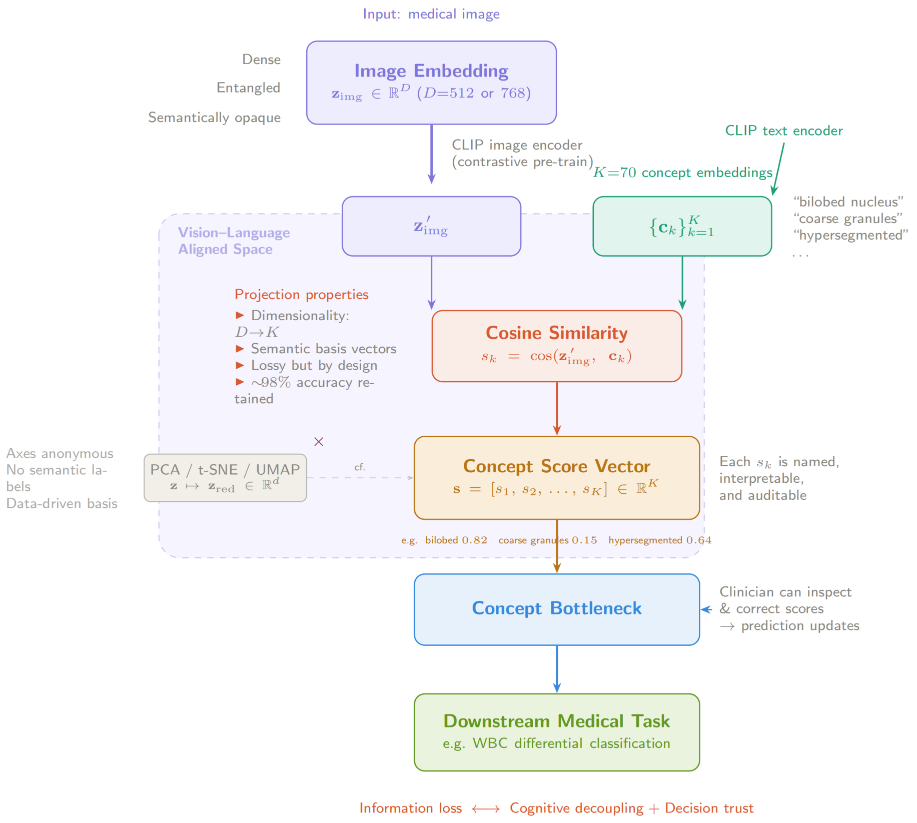


**Figure S3** Cross-modal concept projection pipeline. An image embedding is projected onto $n$ semantically labelled concept axes via cosine similarity in the ConceptCLIP-aligned latent space, yielding an interpretable concept score vector that feeds into the Soft Decision Tree for classification. Unlike classical dimensionality reduction (PCA, t-SNE, UMAP), the projection axes are specified by human domain knowledge encoded as text embeddings rather than derived from data statistics.

### Supplementary Tables: Concept Set Compositions

#### Supplementary Table S1: Full 70-Concept Listing

The following table lists all 70 concepts in the selected concept set, organised by the five morphological categories described in the Methods section. This concept set was used for all main experiments.

Table S1: Complete listing of the 70-concept set.

| # | Concept | Category |
|---|---|---|
| 1 | Segmented nucleus | Nuclear morphology and N:C ratio |
| 2 | Band nucleus (band form) | Nuclear morphology and N:C ratio |
| 3 | Bilobed nucleus | Nuclear morphology and N:C ratio |
| 4 | Round nucleus | Nuclear morphology and N:C ratio |
| 5 | Reniform / indented nucleus | Nuclear morphology and N:C ratio |
| 6 | Coarse chromatin pattern | Nuclear morphology and N:C ratio |
| 7 | Prominent nucleoli | Nuclear morphology and N:C ratio |
| 8 | Irregular nuclear membrane | Nuclear morphology and N:C ratio |
| 9 | Eccentric nucleus | Nuclear morphology and N:C ratio |
| 10 | Binucleated cell | Nuclear morphology and N:C ratio |
| 11 | Multinucleated cell | Nuclear morphology and N:C ratio |
| 12 | Clefted / notched nucleus | Nuclear morphology and N:C ratio |
| 13 | Oval nucleus | Nuclear morphology and N:C ratio |
| 14 | Pyknotic nucleus | Nuclear morphology and N:C ratio |
| 15 | Apoptotic nuclear fragments | Nuclear morphology and N:C ratio |
| 16 | Hypersegmented nucleus | Nuclear morphology and N:C ratio |
| 17 | Fine/open chromatin pattern | Nuclear morphology and N:C ratio |
| 18 | High nuclear-to-cytoplasmic ratio | Nuclear morphology and N:C ratio |
| 19 | Nuclear blebs / micronuclei | Nuclear morphology and N:C ratio |
| 20 | Marked nuclear size variation (anisokaryosis) | Nuclear morphology and N:C ratio |
| 21 | Basophilic cytoplasm | Cytoplasmic tone and texture |
| 22 | Pale cytoplasm | Cytoplasmic tone and texture |
| 23 | Cytoplasmic vacuoles | Cytoplasmic tone and texture |
| 24 | Ground-glass cytoplasm | Cytoplasmic tone and texture |
| 25 | Foamy cytoplasm | Cytoplasmic tone and texture |
| 26 | Cytoplasmic blebs / projections | Cytoplasmic tone and texture |
| 27 | Perinuclear hof (Golgi zone clearing) | Cytoplasmic tone and texture |
| 28 | Polarized cytoplasm (one-sided) | Cytoplasmic tone and texture |
| 29 | Homogeneous dense cytoplasm | Cytoplasmic tone and texture |
| 30 | Paranuclear cytoplasmic vacuoles | Cytoplasmic tone and texture |
| 31 | Clear cytoplasm | Cytoplasmic tone and texture |
| 32 | Plasmacytoid cytoplasm | Cytoplasmic tone and texture |
| 33 | Peripheral basophilic rim | Cytoplasmic tone and texture |
| 34 | Fine azurophilic granules | Cytoplasmic granules and inclusions |
| 35 | Eosinophilic granules | Cytoplasmic granules and inclusions |
| 36 | Basophilic granules | Cytoplasmic granules and inclusions |
| 37 | Toxic granulation | Cytoplasmic granules and inclusions |
| 38 | Döhle bodies | Cytoplasmic granules and inclusions |
| 39 | Auer rods | Cytoplasmic granules and inclusions |
| 40 | Granules obscure nucleus | Cytoplasmic granules and inclusions |
| 41 | Hypogranular / agranular cytoplasm | Cytoplasmic granules and inclusions |
| 42 | Coarse cytoplasmic granules | Cytoplasmic granules and inclusions |
| 43 | Giant cytoplasmic granules | Cytoplasmic granules and inclusions |
| 44 | Azurophilic granules in lymphocytes | Cytoplasmic granules and inclusions |
| 45 | Cytoplasmic crystalline inclusions | Cytoplasmic granules and inclusions |
| 46 | Cytoplasmic pigment granules | Cytoplasmic granules and inclusions |
| 47 | Phagocytosed cellular debris | Cytoplasmic granules and inclusions |
| 48 | Basophilic stippling of erythrocytes | Cytoplasmic granules and inclusions |

Continued on next page

Table S1 (continued)

| # | Concept | Category |
|---|---|---|
| 49 | Howell–Jolly bodies in erythrocytes | Cytoplasmic granules and inclusions |
| 50 | Pappenheimer bodies in erythrocytes | Cytoplasmic granules and inclusions |
| 51 | Uneven granule size (anisogranularity) | Cytoplasmic granules and inclusions |
| 52 | Polar aggregation of cytoplasmic granules | Cytoplasmic granules and inclusions |
| 53 | Perinuclear sparing of cytoplasmic granules | Cytoplasmic granules and inclusions |
| 54 | Nucleated erythrocyte (erythroblast) | Non-leukocyte blood elements |
| 55 | Platelet fragments / clumps | Non-leukocyte blood elements |
| 56 | Red blood cell central pallor | Non-leukocyte blood elements |
| 57 | Red blood cell fragments (schistocytes) | Non-leukocyte blood elements |
| 58 | Spherocytic red blood cell | Non-leukocyte blood elements |
| 59 | Target red blood cell (codocyte) | Non-leukocyte blood elements |
| 60 | Teardrop red blood cell (dacrocyte) | Non-leukocyte blood elements |
| 61 | Red blood cell anisocytosis (size variation) | Non-leukocyte blood elements |
| 62 | Red blood cell poikilocytosis (shape variation) | Non-leukocyte blood elements |
| 63 | Microcytic red blood cells | Non-leukocyte blood elements |
| 64 | Macrocytic red blood cells | Non-leukocyte blood elements |
| 65 | Rouleaux formation of red blood cells | Non-leukocyte blood elements |
| 66 | Giant platelets | Non-leukocyte blood elements |
| 67 | Stain precipitate (artifact) | Preparation and technical artifacts |
| 68 | Overlapping cell clumps (artifact) | Preparation and technical artifacts |
| 69 | Out-of-focus (artifact) | Preparation and technical artifacts |
| 70 | Motion blur (artifact) | Preparation and technical artifacts |

#### Supplementary Table S2: Concept Compositions Across Granularity Levels

The following tables show which concepts are included at each of the four granularity levels ($n = 30, 50, 70, 90$) evaluated in the granularity ablation study (Table 1 of the main text). Concepts are organised by category; a checkmark (✓) indicates inclusion at the corresponding level. This supports reproducibility of the granularity ablation experiment.

**Table S2.** Concept counts per category at each granularity level.

| Category | $n = 30$ | $n = 50$ | $n = 70$ | $n = 90$ |
|---|---|---|---|---|
| Nuclear morphology and N:C ratio | 8 | 15 | 20 | 22 |
| Cytoplasmic tone and texture | 7 | 10 | 13 | 16 |
| Cytoplasmic granules and inclusions | 8 | 14 | 20 | 25 |
| Non-leukocyte blood elements | 3 | 7 | 13 | 23 |
| Preparation and technical artifacts | 4 | 4 | 4 | 4 |
| **Total** | **30** | **50** | **70** | **90** |

**Table S3.** Nuclear morphology and N:C ratio concepts at each granularity level.

| # | Concept | $n$=30 | $n$=50 | $n$=70 | $n$=90 |
|---|---|---|---|---|---|
| 1 | Segmented nucleus | ✓ | ✓ | ✓ | ✓ |
| 2 | Band nucleus (band form) | ✓ | ✓ | ✓ | ✓ |
| 3 | Bilobed nucleus | ✓ | ✓ | ✓ | ✓ |
| 4 | Round nucleus | ✓ | ✓ | ✓ | ✓ |
| 5 | Reniform / indented nucleus | ✓ | ✓ | ✓ | ✓ |
| 6 | Coarse chromatin pattern | ✓ | ✓ | ✓ | ✓ |
| 7 | Prominent nucleoli | ✓ | ✓ | ✓ | ✓ |
| 8 | Irregular nuclear membrane | ✓ | ✓ | ✓ | ✓ |
| 9 | Eccentric nucleus | | ✓ | ✓ | ✓ |
| 10 | Binucleated cell | | ✓ | ✓ | ✓ |
| 11 | Multinucleated cell | | ✓ | ✓ | ✓ |
| 12 | Clefted / notched nucleus | | ✓ | ✓ | ✓ |
| 13 | Oval nucleus | | ✓ | ✓ | ✓ |
| 14 | Pyknotic nucleus | | ✓ | ✓ | ✓ |
| 15 | Apoptotic nuclear fragments | | ✓ | ✓ | ✓ |
| 16 | Hypersegmented nucleus | | | ✓ | ✓ |
| 17 | Fine/open chromatin pattern | | | ✓ | ✓ |
| 18 | High nuclear-to-cytoplasmic ratio | | | ✓ | ✓ |
| 19 | Nuclear blebs / micronuclei | | | ✓ | ✓ |
| 20 | Marked nuclear size variation (anisokaryosis) | | | ✓ | ✓ |
| 21 | Convoluted / folded nucleus | | | | ✓ |
| 22 | Peripheral chromatin condensation (chromatin margination) | | | | ✓ |

**Table S4.** Cytoplasmic tone and texture concepts at each granularity level.

| # | Concept | $n$=30 | $n$=50 | $n$=70 | $n$=90 |
|---|---|---|---|---|---|
| 1 | Basophilic cytoplasm | ✓ | ✓ | ✓ | ✓ |
| 2 | Pale cytoplasm | ✓ | ✓ | ✓ | ✓ |
| 3 | Cytoplasmic vacuoles | ✓ | ✓ | ✓ | ✓ |
| 4 | Ground-glass cytoplasm | ✓ | ✓ | ✓ | ✓ |
| 5 | Foamy cytoplasm | ✓ | ✓ | ✓ | ✓ |
| 6 | Cytoplasmic blebs / projections | ✓ | ✓ | ✓ | ✓ |
| 7 | Perinuclear hof (Golgi zone clearing) | ✓ | ✓ | ✓ | ✓ |
| 8 | Polarized cytoplasm (one-sided) | | ✓ | ✓ | ✓ |
| 9 | Homogeneous dense cytoplasm | | ✓ | ✓ | ✓ |
| 10 | Paranuclear cytoplasmic vacuoles | | ✓ | ✓ | ✓ |
| 11 | Clear cytoplasm | | | ✓ | ✓ |
| 12 | Plasmacytoid cytoplasm | | | ✓ | ✓ |
| 13 | Peripheral basophilic rim | | | ✓ | ✓ |
| 14 | Cytoplasmic streaming / tails | | | | ✓ |
| 15 | Reticular / lace-like cytoplasm | | | | ✓ |
| 16 | Patchy basophilic cytoplasm | | | | ✓ |

**Table S5.** Cytoplasmic granules and inclusions concepts at each granularity level.

| # | Concept | $n$=30 | $n$=50 | $n$=70 | $n$=90 |
|---|---|---|---|---|---|
| 1 | Fine azurophilic granules | ✓ | ✓ | ✓ | ✓ |
| 2 | Eosinophilic granules | ✓ | ✓ | ✓ | ✓ |
| 3 | Basophilic granules | ✓ | ✓ | ✓ | ✓ |
| 4 | Toxic granulation | ✓ | ✓ | ✓ | ✓ |
| 5 | Döhle bodies | ✓ | ✓ | ✓ | ✓ |
| 6 | Auer rods | ✓ | ✓ | ✓ | ✓ |
| 7 | Granules obscure nucleus | ✓ | ✓ | ✓ | ✓ |
| 8 | Hypogranular / agranular cytoplasm | ✓ | ✓ | ✓ | ✓ |
| 9 | Coarse cytoplasmic granules | | ✓ | ✓ | ✓ |
| 10 | Giant cytoplasmic granules | | ✓ | ✓ | ✓ |
| 11 | Azurophilic granules in lymphocytes | | ✓ | ✓ | ✓ |
| 12 | Cytoplasmic crystalline inclusions | | ✓ | ✓ | ✓ |
| 13 | Cytoplasmic pigment granules | | ✓ | ✓ | ✓ |
| 14 | Phagocytosed cellular debris | | ✓ | ✓ | ✓ |
| 15 | Basophilic stippling of erythrocytes | | | ✓ | ✓ |
| 16 | Howell–Jolly bodies in erythrocytes | | | ✓ | ✓ |
| 17 | Pappenheimer bodies in erythrocytes | | | ✓ | ✓ |
| 18 | Uneven granule size (anisogranularity) | | | ✓ | ✓ |
| 19 | Polar aggregation of cytoplasmic granules | | | ✓ | ✓ |
| 20 | Perinuclear sparing of cytoplasmic granules | | | ✓ | ✓ |
| 21 | Uniform fine cytoplasmic granules | | | | ✓ |
| 22 | Peripheral rim of cytoplasmic granules | | | | ✓ |
| 23 | Perinuclear ring of cytoplasmic granules | | | | ✓ |
| 24 | Discrete non-granular cytoplasmic inclusions | | | | ✓ |
| 25 | Single large cytoplasmic inclusion | | | | ✓ |

**Table S6.** Non-leukocyte blood elements concepts at each granularity level.

| # | Concept | $n$=30 | $n$=50 | $n$=70 | $n$=90 |
|---|---|---|---|---|---|
| 1 | Nucleated erythrocyte (erythroblast) | ✓ | ✓ | ✓ | ✓ |
| 2 | Platelet fragments / clumps | ✓ | ✓ | ✓ | ✓ |
| 3 | Red blood cell central pallor | ✓ | ✓ | ✓ | ✓ |
| 4 | Red blood cell fragments (schistocytes) | | ✓ | ✓ | ✓ |
| 5 | Spherocytic red blood cell | | ✓ | ✓ | ✓ |
| 6 | Target red blood cell (codocyte) | | ✓ | ✓ | ✓ |
| 7 | Teardrop red blood cell (dacrocyte) | | ✓ | ✓ | ✓ |
| 8 | Red blood cell anisocytosis (size variation) | | | ✓ | ✓ |
| 9 | Red blood cell poikilocytosis (shape variation) | | | ✓ | ✓ |
| 10 | Microcytic red blood cells | | | ✓ | ✓ |
| 11 | Macrocytic red blood cells | | | ✓ | ✓ |
| 12 | Rouleaux formation of red blood cells | | | ✓ | ✓ |
| 13 | Giant platelets | | | ✓ | ✓ |
| 14 | Oval / elliptic red blood cell (ovalocyte / elliptocyte) | | | | ✓ |
| 15 | Stomatocytic red blood cell (stomatocyte) | | | | ✓ |
| 16 | Echinocytic red blood cell (burr cell) | | | | ✓ |
| 17 | Acanthocytic red blood cell (spur cell) | | | | ✓ |
| 18 | Hypochromic red blood cell | | | | ✓ |
| 19 | Polychromatophilic red blood cell | | | | ✓ |
| 20 | Platelet anisocytosis (size variation) | | | | ✓ |
| 21 | Hypogranular platelets | | | | ✓ |
| 22 | Hypergranular platelets | | | | ✓ |
| 23 | Platelet satellitism around leukocytes | | | | ✓ |

**Table S7.** Preparation and technical artifacts concepts. All four concepts are shared across all granularity levels.

| # | Concept | $n$=30 | $n$=50 | $n$=70 | $n$=90 |
|---|---|---|---|---|---|
| 1 | Stain precipitate (artifact) | ✓ | ✓ | ✓ | ✓ |
| 2 | Overlapping cell clumps (artifact) | ✓ | ✓ | ✓ | ✓ |
| 3 | Out-of-focus (artifact) | ✓ | ✓ | ✓ | ✓ |
| 4 | Motion blur (artifact) | ✓ | ✓ | ✓ | ✓ |

### Supplementary Note S1: Complete SDT Node Annotations

This note provides detailed histological annotations for the 15 internal nodes of the trained depth-4 Soft Decision Tree (SDT), complementing Figure 3 and Figure 4 of the main text. The overall tree structure is shown in Supplementary Figure S1. At each internal node, samples with high decision scores $\mathbf{w}^\top \mathbf{c} + b$ are routed left; samples with low scores are routed right.

*Conventions.*
For nodes with sufficient statistical support (≥50 samples; see Methods, Pruning and Tree Annotation), representative images are drawn from the Top-50 (highest decision score) and Bottom-50 (lowest decision score) training samples routed to that node. For structurally necessary but informationally redundant nodes, a brief functional description replaces image-based annotation. Pruned nodes (fewer than 50 samples or both children pruned) are listed only in the overview tables below.

#### Overview Tables

**Table S8.** Summary of all 15 internal nodes.

| Node | Layer | Samples | Status | Left child ($n$) | Right child ($n$) | Primary discrimination |
|---|---|---|---|---|---|---|
| IN 0 | 1 | 13,671 | Active | IN 1 (5,204) | IN 2 (8,467) | Cytoplasmic staining tonality |
| IN 1 | 2 | 5,204 | Active | IN 3 (4,146) | IN 4 (1,058) | Lymphocyte isolation |
| IN 2 | 2 | 8,467 | Active | IN 5 (1,502) | IN 6 (6,965) | Erythroblast isolation |
| IN 3 | 3 | 4,146 | Active | IN 7 (1,020) | IN 8 (3,126) | Basophil vs. IG/monocyte |
| IN 4 | 3 | 1,058 | Active | IN 9 (1,057) | IN 10 (1) | Pass-through (lymphocyte) |
| IN 5 | 3 | 1,502 | Active | IN 11 (1,479) | IN 12 (23) | Pre-filter for erythroblast/IG |
| IN 6 | 3 | 6,965 | Active | IN 13 (5,105) | IN 14 (1,860) | Granulocyte vs. platelet |
| IN 7 | 4 | 1,020 | Active | L0 (986) | L1 (34) | Basophil consolidation |
| IN 8 | 4 | 3,126 | Active | L2 (1,961) | L3 (1,165) | IG (promyelocyte) vs. monocyte |
| IN 9 | 4 | 1,057 | Active | L4 (1,004) | L5 (53) | Lymphocyte consolidation |
| IN 10 | 4 | 1 | **Pruned** | L6 (1) | L7 (0) | — |
| IN 11 | 4 | 1,479 | Active | L8 (1,156) | L9 (323) | Erythroblast vs. IG (metamyelocyte) |
| IN 12 | 4 | 23 | **Pruned** | L10 (16) | L11 (7) | — |
| IN 13 | 4 | 5,105 | Active | L12 (2,627) | L13 (2,478) | Neutrophil vs. eosinophil |
| IN 14 | 4 | 1,860 | Active | L14 (1,831) | L15 (29) | Platelet consolidation |

**Table S9.** Summary of all 16 leaf nodes.

| Leaf | Predicted class | Confidence | Samples | Status |
|---|---|---|---|---|
| L0 | Basophil | 99% | 986 | Active |
| L1 | Eosinophil | 26% | 34 | **Pruned** |
| L2 | Immature granulocyte | 99% | 1,961 | Active |
| L3 | Monocyte | 99% | 1,165 | Active |
| L4 | Lymphocyte | 99% | 1,004 | Active |
| L5 | Platelet | 41% | 53 | **Pruned** |
| L6 | — | — | 1 | **Pruned** |
| L7 | — | — | 0 | **Pruned** |
| L8 | Erythroblast | 99% | 1,156 | Active |
| L9 | Immature granulocyte | 86% | 323 | Active |
| L10 | Platelet | 27% | 16 | **Pruned** |
| L11 | Lymphocyte | 26% | 7 | **Pruned** |
| L12 | Neutrophil | 99% | 2,627 | Active |
| L13 | Eosinophil | 99% | 2,478 | Active |
| L14 | Platelet | 99% | 1,831 | Active |
| L15 | Monocyte | 21% | 29 | **Pruned** |

### IN 0 — Layer 1 (Root) — 13,671 samples

**Routing:** All training samples. Left child: IN 1 (5,204 samples). Right child: IN 2 (8,467 samples).

#### Left branch (Top-50, high routing score)

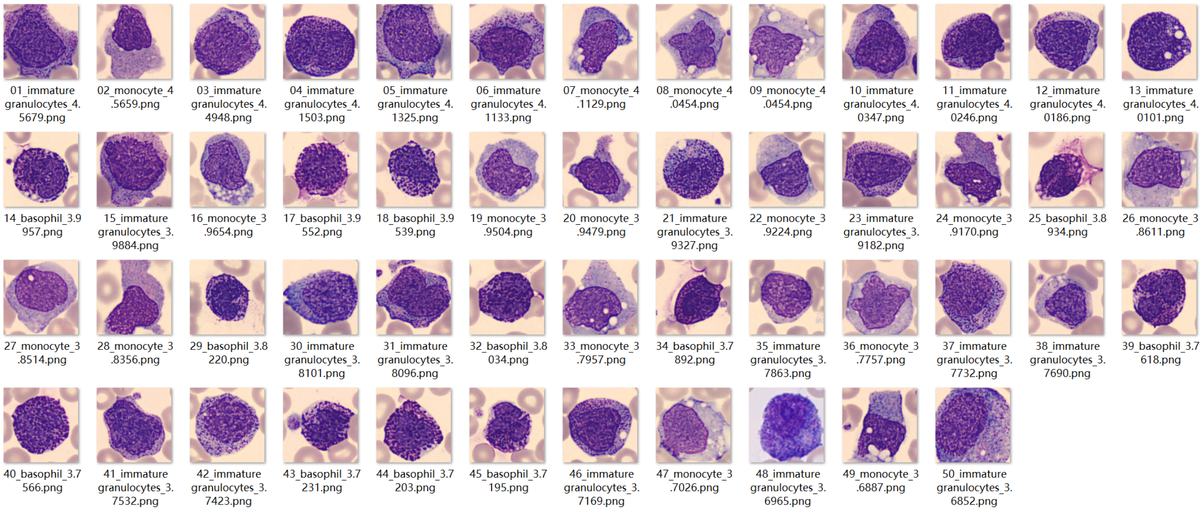


**Dominant class in Top-50:** Immature granulocyte (22/50). Also represented: monocytes and basophils.

**Histological impression.** High-score samples are heterogeneous, comprising immature granulocytes, monocytes, and basophils—cell types that share a common gross appearance under May–Grünwald–Giemsa staining: cytoplasm tending toward blue or pale violet tones (basophilic staining character). The root node filter primarily separates cells by overall cytoplasmic staining tonality rather than nuclear morphology.

#### Right branch (Bottom-50, low routing score)

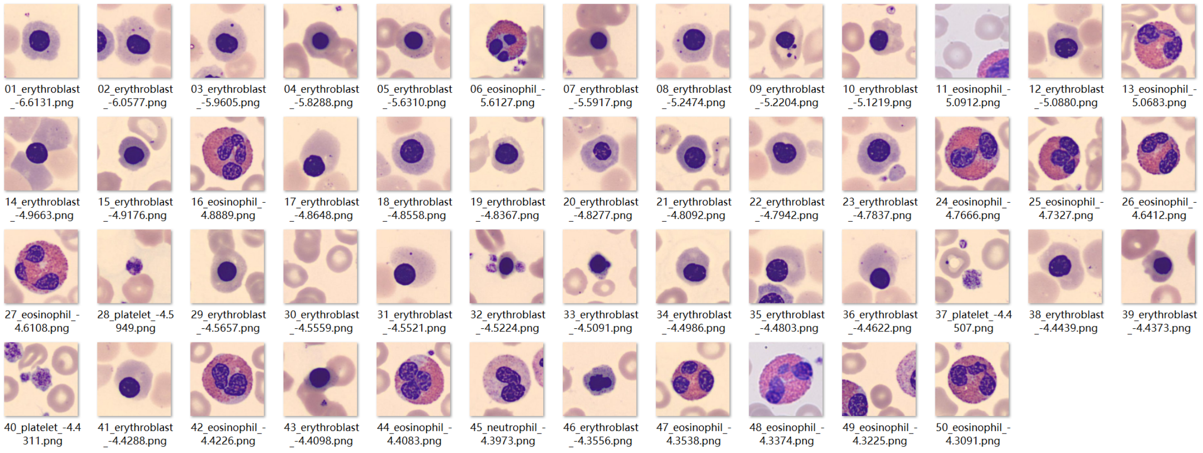


**Dominant class in Bottom-50:** Erythroblast (32/50). Also represented: eosinophils, neutrophils, and platelets.

**Histological impression.** Low-score samples tend toward pink-to-red cytoplasmic staining (eosinophilic character) or small cell body size (platelets), contrasting with the basophilic-toned cells routed left.

**Remark.** The root split functions as a coarse chromatic separator, partitioning the eight-class problem into two morphologically coherent subsets. This global division does not yet resolve individual cell types but establishes the primary organising axis of routine blood smear analysis.

### IN 1 — Layer 2 — 5,204 samples (left subtree of IN 0)

**Routing:** Basophilic-toned cells from IN 0. Left child: IN 3 (4,146 samples). Right child: IN 4 (1,058 samples).

#### Left branch (Top-50, high routing score)

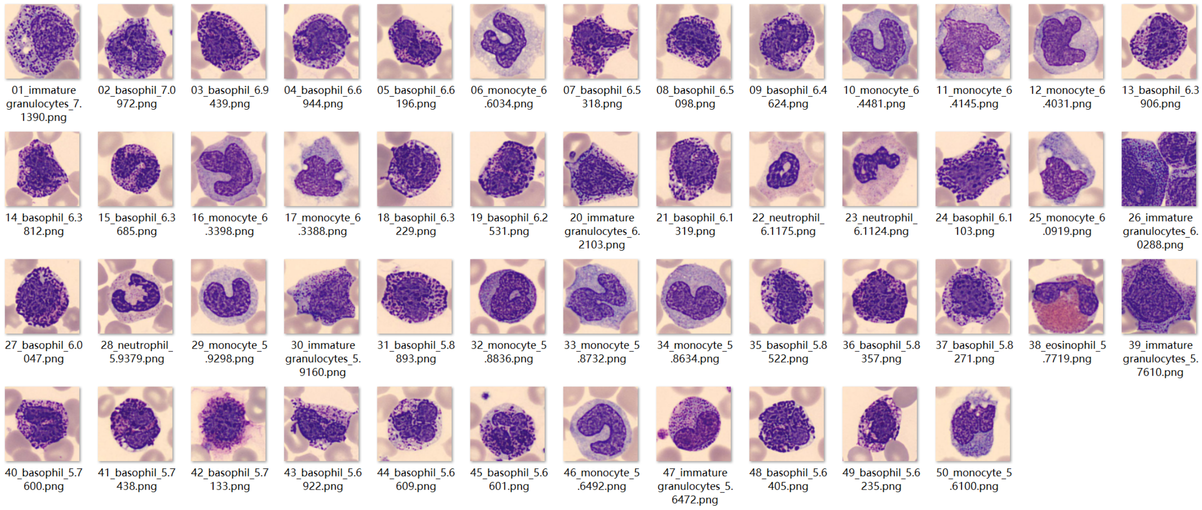


**Dominant class in Top-50:** Basophil (27/50). Also represented: immature granulocytes and monocytes.

**Histological impression.** High-score samples comprise basophils, immature granulocytes, and monocytes—cell types sharing relatively large cell bodies, irregularly shaped nuclei (reniform or horseshoe-shaped), and cytoplasm with an overall basophilic tone containing azurophilic granules.

#### Right branch (Bottom-50, low routing score)

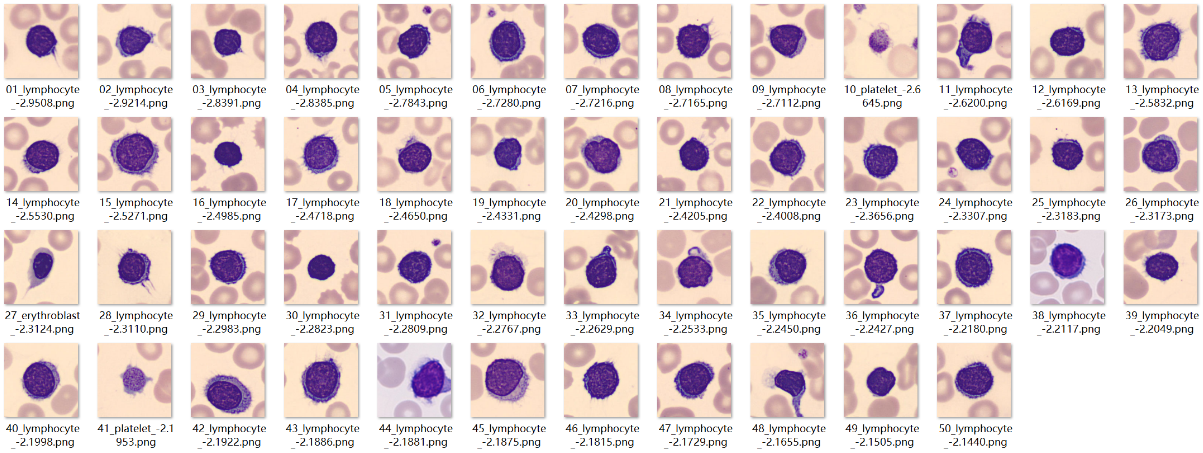


**Dominant class in Bottom-50:** Lymphocyte (47/50).

**Histological impression.** Low-score samples are predominantly lymphocytes. Lymphocytes are morphologically distinct from the other cell types in this subtree: small cell body, very high N/C ratio, a round and densely chromatic nucleus occupying nearly the entire cell, and only a thin peripheral rim of basophilic cytoplasm without visible granules. The filter at IN 1 effectively separates lymphocytes from the larger, granule-bearing cell types.

**Remark.** The discriminative features used by IN 1—small cell body, high N/C ratio, and dense round nucleus—are precisely the standard textbook criteria for lymphocyte identification in peripheral blood smear analysis.

### IN 2 — Layer 2 — 8,467 samples (right subtree of IN 0)

**Routing:** Eosinophilic-toned and small cells from IN 0. Left child: IN 5 (1,502 samples). Right child: IN 6 (6,965 samples).

#### Left branch (Top-50, high routing score)

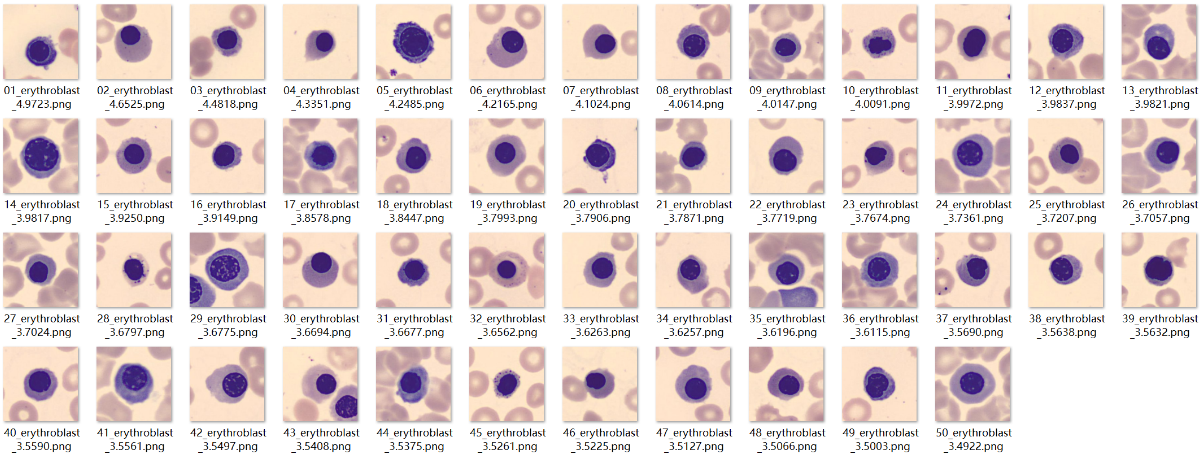


**Dominant class in Top-50:** Erythroblast (50/50).

**Histological impression.** All 50 high-score samples are erythroblasts (nucleated red blood cell precursors). Their defining features include a small, round, densely stained nucleus (dark blue/purple) and cytoplasm ranging from basophilic (blue) to polychromatic (grey-pink) depending on the maturation stage. The filter at IN 2 isolates erythroblasts with high selectivity.

#### Right branch (Bottom-50, low routing score)

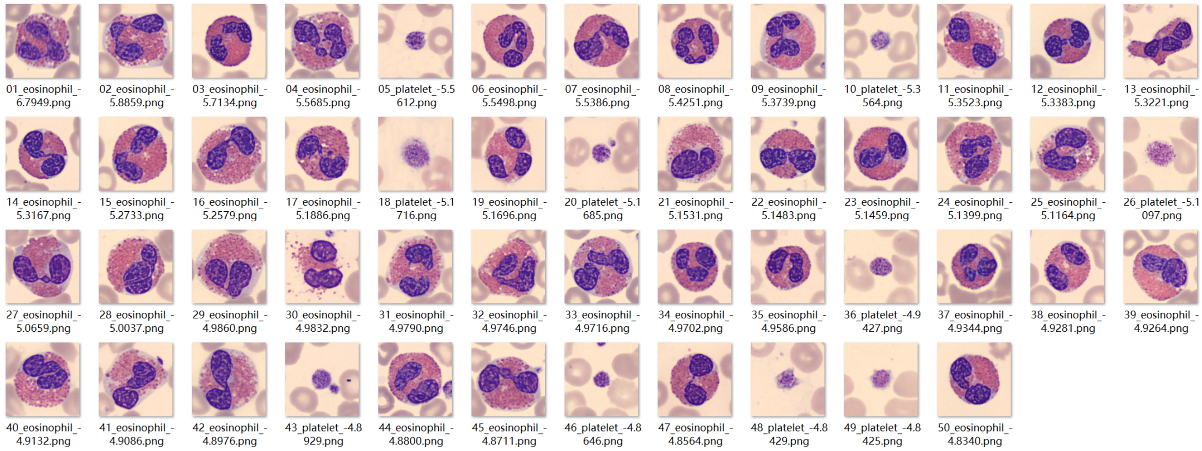


**Dominant class in Bottom-50:** Eosinophil (40/50). Also represented: platelets.

**Histological impression.** Low-score samples are predominantly eosinophils, with platelets also present—cell types sharing eosinophilic cytoplasmic staining and, in eosinophils, lobated nuclear morphology. These features contrast with the round-nucleated, basophilic-to-polychromatic erythroblasts routed left.

**Remark.** IN 2 provides an early separation of erythroblasts from the remaining eosinophilic-toned population, consistent with the clinical morphological criterion that distinguishes nucleated red cell precursors by their round, pyknotic nucleus and shifting cytoplasmic basophilia.

### IN 3 — Layer 3 — 4,146 samples (left branch of IN 1)

**Routing:** Basophils, immature granulocytes, and monocytes from IN 1. Left child: IN 7 (1,020 samples). Right child: IN 8 (3,126 samples).

#### Left branch (Top-50, high routing score)

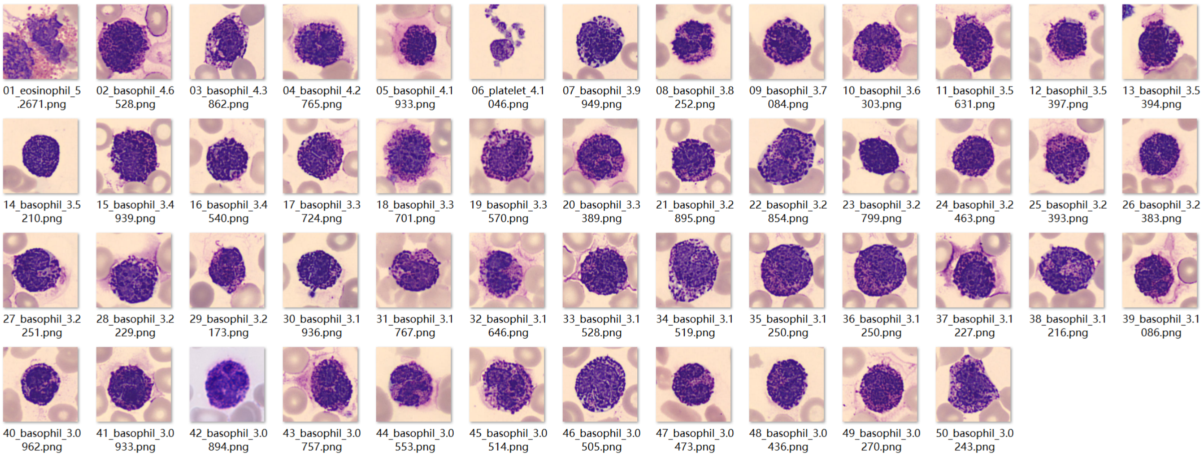


**Dominant class in Top-50:** Basophil (48/50).

**Histological impression.** High-score samples are nearly exclusively basophils. The cytoplasm is packed with coarse, irregularly sized, deep purple-black basophilic granules that are so dense they frequently obscure the underlying nucleus—the hallmark "obscuring granules" pattern. The nucleus, though partially hidden, is typically bilobed or trilobed.

#### Right branch (Bottom-50, low routing score)

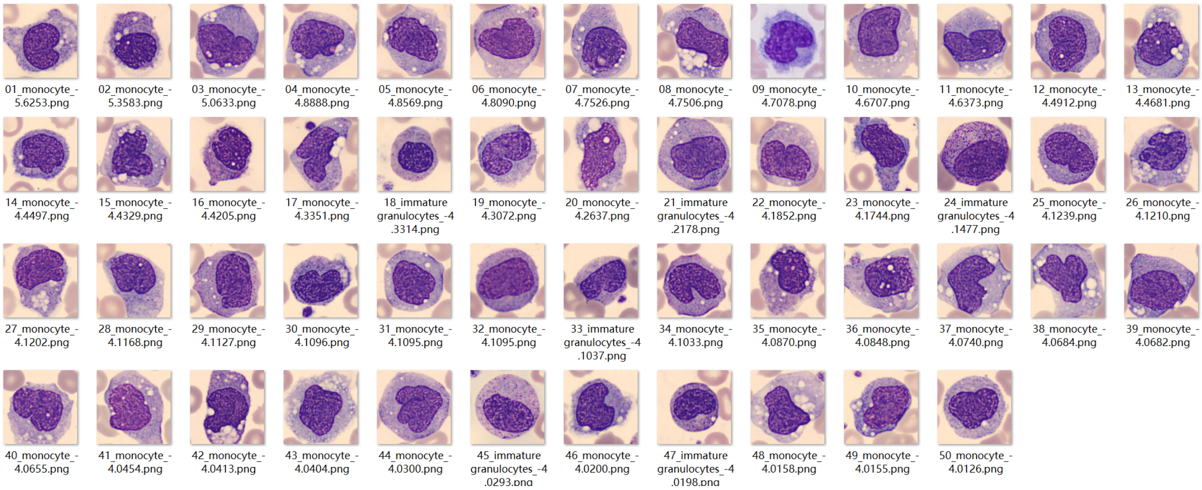


**Dominant class in Bottom-50:** Monocyte (44/50). Also represented: immature granulocytes.

**Histological impression.** Low-score samples are dominated by monocytes and immature granulocytes. Common features include irregular cell margins, a hazy "ground-glass" cytoplasmic appearance, and irregularly shaped nuclei. Unlike the basophils routed left, neither cell type exhibits the dense nucleus-obscuring granulation pattern.

**Remark.** IN 3 addresses a morphologically challenging distinction: basophils versus immature granulocytes (particularly promyelocytes), both of which contain dense purple-red granules under routine staining. The filter successfully leverages whether the granules completely obscure the nucleus (basophil) or leave the nuclear boundary discernible (IG/monocyte).

### IN 6 — Layer 3 — 6,965 samples (right branch of IN 2)

**Routing:** Eosinophilic-toned non-erythroblast cells from IN 2. Left child: IN 13 (5,105 samples). Right child: IN 14 (1,860 samples).

#### Left branch (Top-50, high routing score)

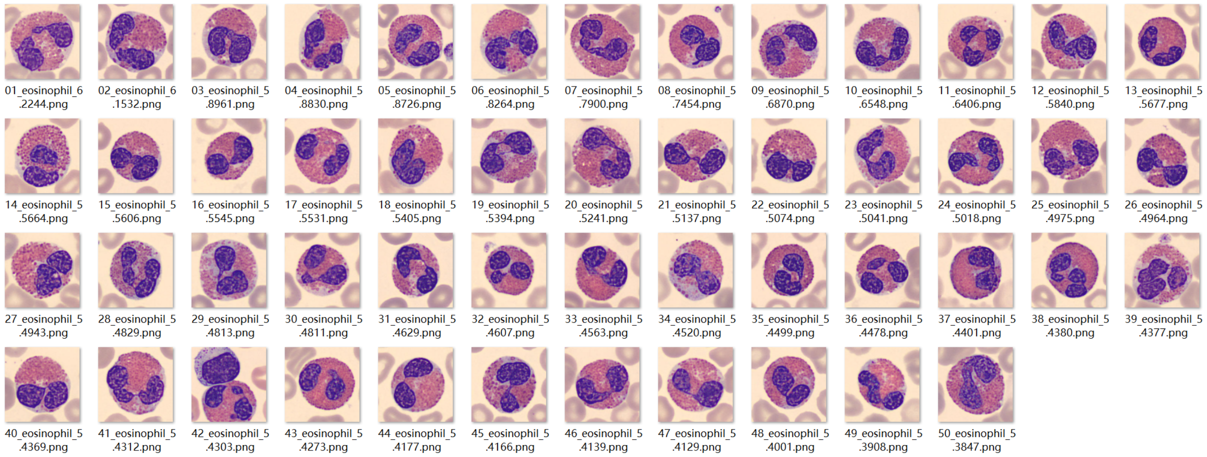


**Dominant class in Top-50:** Eosinophil (50/50).

**Histological impression.** All 50 high-score samples are eosinophils. The filter captures their defining features: cytoplasm densely packed with coarse, uniform, brick-red granules exhibiting characteristic refractile quality, and a typical bilobed nucleus stained purple.

#### Right branch (Bottom-50, low routing score)

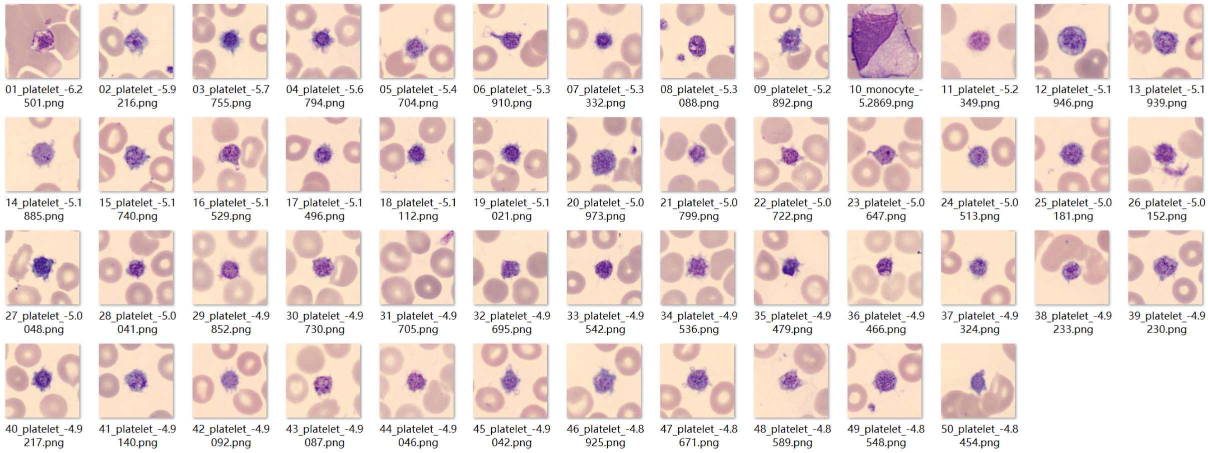


**Dominant class in Bottom-50:** Platelet (49/50).

**Histological impression.** Low-score samples are nearly exclusively platelets: anucleate, small, appearing as purple-red granular bodies with indistinct margins. Their minimal cell body size and absence of a nucleus clearly distinguish them from the large, granule-filled eosinophils routed left.

**Remark.** IN 6 performs the first separation between nucleated granulocytes (eosinophils and neutrophils, routed left) and platelets (routed right). The downstream node IN 13 then resolves the eosinophil–neutrophil distinction.

### IN 8 — Layer 4 — 3,126 samples (right branch of IN 3)

**Routing:** Immature granulocytes and monocytes remaining after IN 3 routes basophils left. Left child: L2 (1,961 samples). Right child: L3 (1,165 samples).

#### Left branch (Top-50, high routing score) → Leaf L2

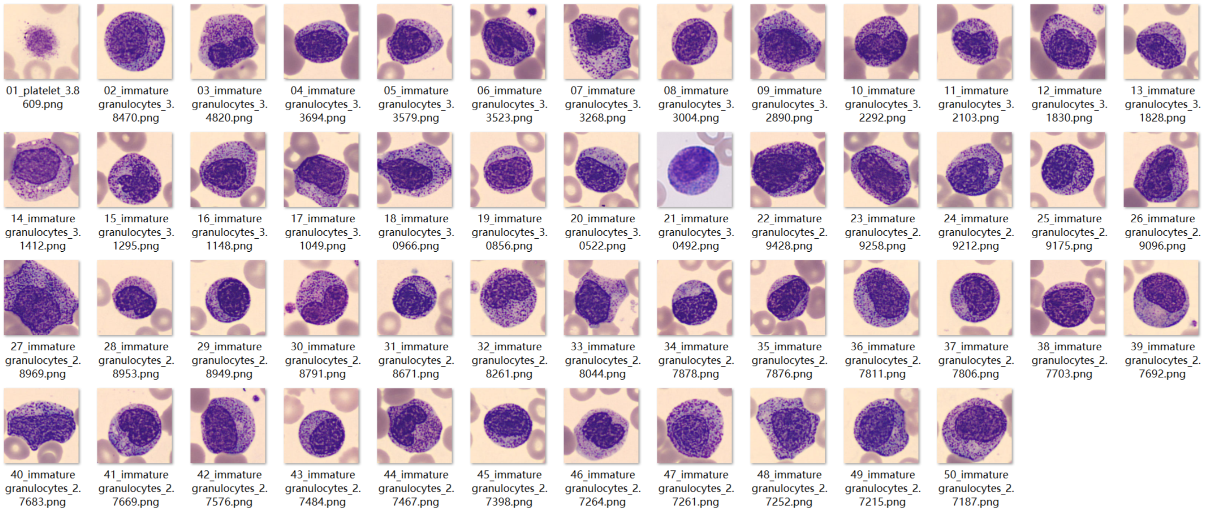


**Dominant class in Top-50:** Immature granulocyte (49/50).

**Histological impression.** High-score samples display typical immature granulocyte morphology at the promyelocyte stage: round-to-oval nuclei without indentation, dense azurophilic (primary) granules that, unlike in basophils, do not completely obscure the nuclear boundary, and a relatively high N/C ratio.

#### Right branch (Bottom-50, low routing score) → Leaf L3

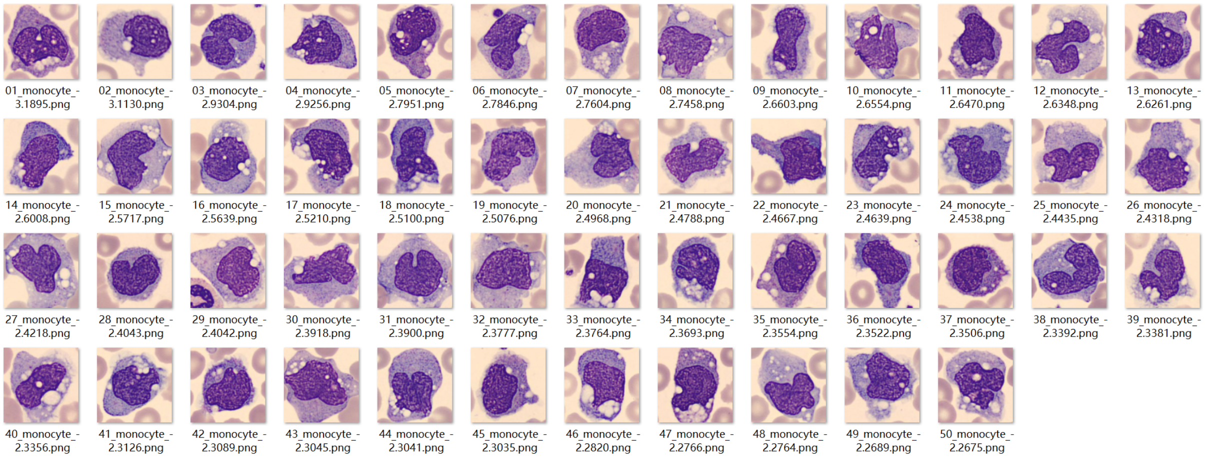


**Dominant class in Bottom-50:** Monocyte (50/50).

**Histological impression.** Low-score samples are uniformly monocytes: irregular cell margins with occasional pseudopodia, diffuse "ground-glass" cytoplasm due to fine unresolvable azurophilic granules (lysosomes), scattered grey-white vacuoles, and a reniform or horseshoe-shaped nucleus.

**Leaf outcomes:** L2 — immature granulocyte, 99% confidence, 1,961 samples (active). L3 — monocyte, 99% confidence, 1,165 samples (active).

**Remark.** Both downstream leaves achieve 99% prediction confidence, indicating a clean separation. The IG samples captured by L2 predominantly exhibit promyelocyte-stage features. A second IG-predicting leaf (L9, via IN 11) captures metamyelocyte-stage features via a different decision path—see IN 11 below for details of this emergent subtype separation.

### IN 11 — Layer 4 — 1,479 samples (left branch of IN 5)

**Routing:** Mixed erythroblasts and immature granulocytes, pre-enriched by IN 2 and IN 5. Left child: L8 (1,156 samples). Right child: L9 (323 samples).

#### Left branch (Top-50, high routing score) → Leaf L8

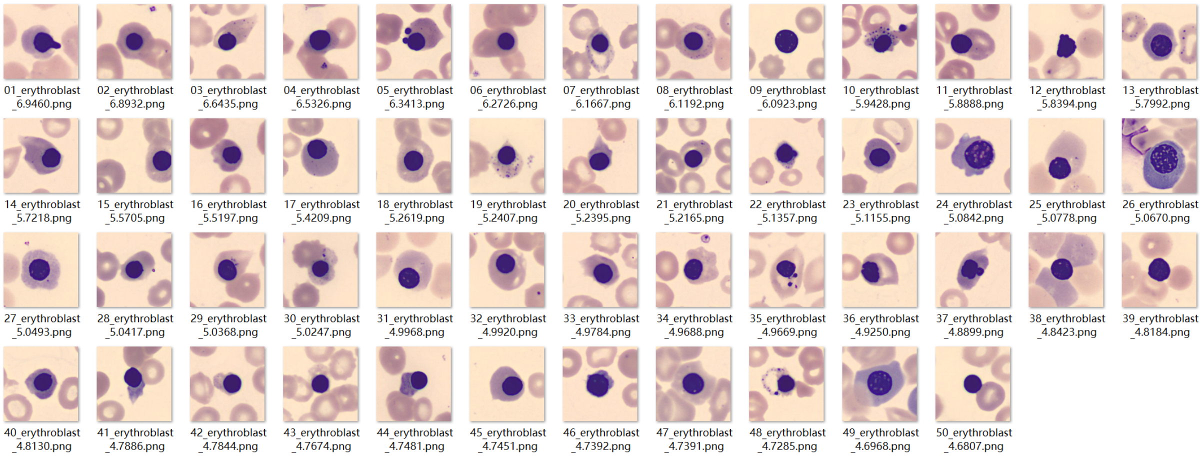


**Dominant class in Top-50:** Erythroblast (50/50).

**Histological impression.** All 50 high-score samples are erythroblasts displaying late-stage morphology: small cell bodies, round nuclei with smooth boundaries and high chromatin density (progressing toward pyknosis), and cytoplasm shifting from basophilic to polychromatic or orthochromatic (pink-red). Occasional serrated cytoplasmic margins likely reflect smear preparation artifacts.

#### Right branch (Bottom-50, low routing score) → Leaf L9

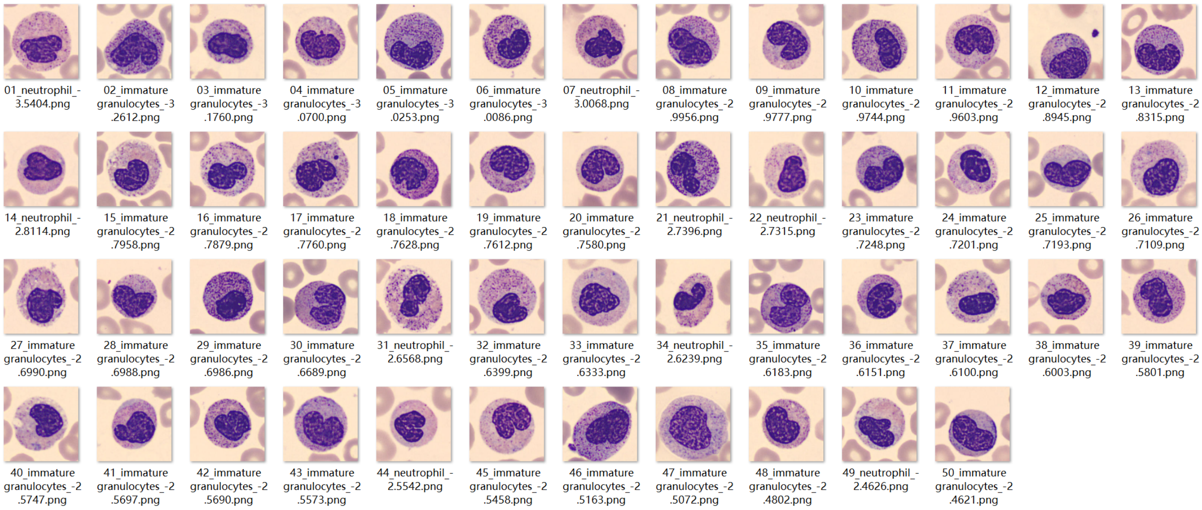


**Dominant class in Bottom-50:** Immature granulocyte (41/50).

**Histological impression.** Low-score samples are predominantly immature granulocytes at the *metamyelocyte* stage: the nucleus shows distinct indentation (reniform or kidney-shaped), the cytoplasm is pale pink approaching the appearance of mature neutrophils, and the N/C ratio is lower than that of promyelocytes. These morphological features contrast sharply with the L2 population (see IN 8), which exhibits promyelocyte-stage characteristics.

**Leaf outcomes:** L8 — erythroblast, 99% confidence, 1,156 samples (active). L9 — immature granulocyte, 86% confidence, 323 samples (active).

**Remark.** IN 11 separates erythroblasts from immature granulocytes within the morphologically overlapping population enriched by the earlier nodes. Critically, the IG samples reaching L9 exhibit metamyelocyte-stage features (indented nucleus, pale cytoplasm, reduced N/C ratio), whereas those at L2 (via IN 8) exhibit promyelocyte-stage features (round nucleus, dense azurophilic granules, high N/C ratio). The SDT has thus learned to separate two morphologically distinct maturation stages of immature granulocytes—promyelocyte and metamyelocyte—along entirely different decision paths, despite the absence of subtype labels in BloodMNIST. This finding is discussed in the main text.

### IN 13 — Layer 4 — 5,105 samples (left branch of IN 6)

**Routing:** Eosinophils and neutrophils from IN 6. Left child: L12 (2,627 samples). Right child: L13 (2,478 samples).

#### Left branch (Top-50, high routing score) → Leaf L12

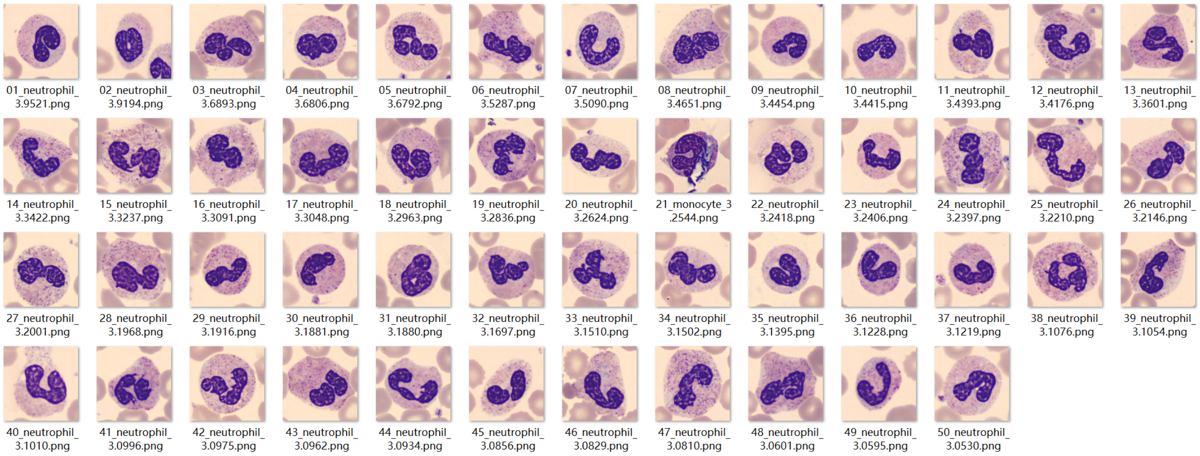


**Dominant class in Top-50:** Neutrophil (49/50).

**Histological impression.** High-score samples are nearly exclusively neutrophils, characterised by a multilobed (typically 3–5 lobes) nucleus with condensed chromatin and cytoplasm containing fine, neutral-staining (pink-purple) specific granules.

#### Right branch (Bottom-50, low routing score) → Leaf L13

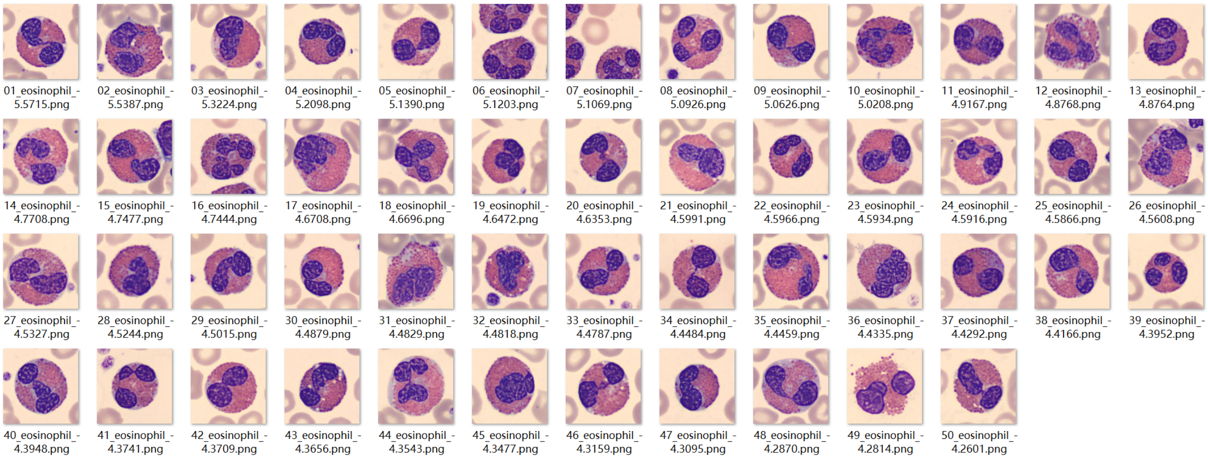


**Dominant class in Bottom-50:** Eosinophil (50/50).

**Histological impression.** All 50 low-score samples are eosinophils: cytoplasm filled with coarse, uniform, brick-red granules with characteristic refractile ("glassy bead") quality and a typically bilobed nucleus—a key differentiator from the more extensively segmented neutrophil nucleus.

**Leaf outcomes:** L12 — neutrophil, 99% confidence, 2,627 samples (active). L13 — eosinophil, 99% confidence, 2,478 samples (active).

**Remark.** IN 13 resolves the neutrophil–eosinophil distinction. The two cell types share lobated nuclear morphology and granule-rich cytoplasm but differ in granule type (fine neutral-staining vs. coarse brick-red) and degree of nuclear segmentation (multi-lobed vs. bilobed). Both downstream leaves achieve 99% prediction confidence.

#### Structurally Necessary Nodes (Brief Annotations)

The following five nodes are informationally redundant (pass-through nodes that do not introduce new discriminative criteria) or serve primarily as consolidation steps. They are annotated briefly for completeness.

**IN 4** (Layer 3, 1,058 samples; parent: IN 1 right branch): Routes 1,057 of 1,058 samples to the left child (IN 9), which leads to the lymphocyte leaf L4. One sample reaches the right child (IN 10, pruned). IN 4 does not introduce a new discriminative split; the lymphocyte population was already effectively isolated at IN 1.

**IN 5** (Layer 3, 1,502 samples; parent: IN 2 left branch): Routes 1,479 of 1,502 samples to the left child (IN 11), which resolves erythroblasts from metamyelocyte-stage IGs. The remaining 23 samples reach IN 12 (pruned). The Top-50 analysis at IN 5 reveals a shared morphological signature between erythroblasts and immature granulocytes (pinkish cytoplasm, purple nuclei, round nuclear contours), establishing the morphological context that IN 11 must then resolve.

**IN 7** (Layer 4, 1,020 samples; parent: IN 3 left branch): Routes 986 of 1,020 samples to L0 (basophil, 99% confidence). The remaining 34 samples reach L1 (pruned, 26% confidence). IN 7 consolidates the basophil population already enriched by IN 3. Decision path IN 0 → IN 1 → IN 3 → IN 7 → L0 constitutes the complete basophil recognition pathway.

**IN 9** (Layer 4, 1,057 samples; parent: IN 4 left branch): Routes 1,004 of 1,057 samples to L4 (lymphocyte, 99% confidence). The remaining 53 samples reach L5 (pruned, 41% confidence). Decision path IN 0 → IN 1 → IN 4 → IN 9 → L4 constitutes the lymphocyte recognition pathway; IN 4 and IN 9 serve as structural consequences of the binary tree constraint.

**IN 14** (Layer 4, 1,860 samples; parent: IN 6 right branch): Routes 1,831 of 1,860 samples to L14 (platelet, 99% confidence). The remaining 29 samples reach L15 (pruned, 21% confidence). Decision path IN 0 → IN 2 → IN 6 → IN 14 → L14 constitutes the platelet recognition pathway.

### Supplementary Note S2: Text Prompt Templates

The following ten prompt templates were used to encode each morphological concept as a text embedding ensemble. For each concept, the placeholder {} is replaced by the concept name (e.g., "Segmented nucleus" or "Basophilic granules"). The ten resulting text embeddings are averaged to produce a single robust representation per concept, mitigating the linguistic bias of any individual phrasing. This ensemble strategy follows standard practice for zero-shot inference with CLIP-based models.

**Table S10.** Text prompt templates used for concept embedding.

| # | Template |
|---|---|
| 1 | "a blood cell photo with sign of {}" |
| 2 | "a photo of a blood cell with {}" |
| 3 | "a blood cell image indicating {}" |
| 4 | "an image of a blood cell showing {}" |
| 5 | "peripheral blood cell with {}" |
| 6 | "a peripheral blood cell photo with sign of {}" |
| 7 | "a photo of a peripheral blood cell with {}" |
| 8 | "a peripheral blood cell image indicating {}" |
| 9 | "an image of blood cell showing {}" |
| 10 | "a micrograph showing the feature of {}" |
